# Determinants of trafficking, conduction, and disease within a K^+^ channel revealed through multiparametric deep mutational scanning

**DOI:** 10.1101/2022.01.06.475280

**Authors:** Willow Coyote-Maestas, David Nedrud, Yungui He, Daniel Schmidt

**Author notes:** Correspondence and requests for materials should be addressed to D.S. Department of Bioengineering and Therapeutic Science and Quantitative Biosciences Institute, University of California, San Francisco, San Francisco, CA, 94158.

## Abstract

A longstanding goal in protein science and clinical genetics is to develop quantitative models of sequence, structure, and function relationships and delineate the mechanisms by which mutations cause disease. Deep Mutational Scanning (DMS) is a promising strategy to map how amino acids contribute to protein structure and function and to advance clinical variant interpretation. Here, we introduce 7,429 single residue missense mutation into the Inward Rectifier K^+^ channel Kir2.1 and determine how this affects folding, assembly, and trafficking, as well as regulation by allosteric ligands and ion conduction. Our data provides high-resolution information on a cotranslationally­folded biogenic unit, trafficking and quality control signals, and segregated roles of different structural elements in fold-stability and function. We show that Kir2.1 trafficking mutants are underrepresented in variant effect databases, which has implications for clinical practice. By comparing fitness scores with expert-reviewed variant effects, we can predict the pathogenicity of ‘variants of unknown significance’ and disease mechanisms of know pathogenic mutations. Our study in Kir2.1 provides a blueprint for how multiparametric DMS can help us understand the mechanistic basis of genetic disorders and the structure-function relationships of proteins.

## Introduction

Inward rectifier K^+^ (Kir) play a central role in physiology by setting, maintaining, and regulating a cell’s resting membrane potential ^1^. Misregulated Kir cause neurological disorders (alcohol and opiate addiction^2–4^, Down’s syndrome ^5^, epilepsy ^6^, Parkinsons ^7^), cardiac disorders (Long QT syndrome ^8^), hereditary renal diseases^9^, and diabetes ^10^.

While some mechanisms that underlie disease-causing Kir variants involve aberrant function (gating, ion permeation, ligand binding ^11^) or transcript processing ^12^, disease-causing variants are often linked to trafficking defects. For example, over half the missense mutations investigated in Kir1.1 (ROMK1) affected surface expression, often caused by proteosomal degradation of folding-deficient or mistrafficked channels ^13, 14^. A comparison of mutation hotspots in the Kir C-terminal domain (CTD) revealed that ∼60% of variants could be linked to impaired surface expression ^15, 16^. Many mutations linked to neonatal diabetes reduced Kir6.2 surface expression to varying degrees ^17^. Beyond known trafficking signals (e.g., ER and Golgi-export signal sequences ^18–23)^, mutations along the entire Kir primary sequence can disrupt surface expression16. Several additional factors control surface expression, such as protein stability, interactions with trafficking partners, and complexes that stabilize channels in the membrane ^20, 24, 25^.

The apparent prevalence of Kir trafficking phenotypes has a significant caveat: phenotypes are experimentally determined for a small fraction of possible or clinically observed variants. Of the 8,113 possible missense mutations for KCNJ2/Kir2.1 only 163 are represented in ClinVar– a database of clinically observed variation and phenotypes ^26^ (**Fig. 1A**, **Supplementary Fig. 1**). Most studies focus on common natural variants observed in human genomes or exomes. Rare genetic variants vastly outnumber common variants ^27^ and are implicated in a substantial portion of complex human disease ^28^. Rarity means that clinical significance (benign or pathogenic) of most variants cannot be established using genome-wide association studies due to low statistical power for calling pathogenic variants. For KCNJ2/Kir2.1 ClinVar reports 144 missense mutations, but most (72%) have “uncertain significance” or conflicting or no interpretation (**Fig. 1A**). Computational algorithms (PolyPhen-2 ^29^, SIFT ^30^, EVE ^31^ etc.) are currently filling this gap by predicting the consequence of mutations. However, computational approaches work best in conserved regions of genes and cannot predict mechanism of pathogenicity.

**Figure 1.**
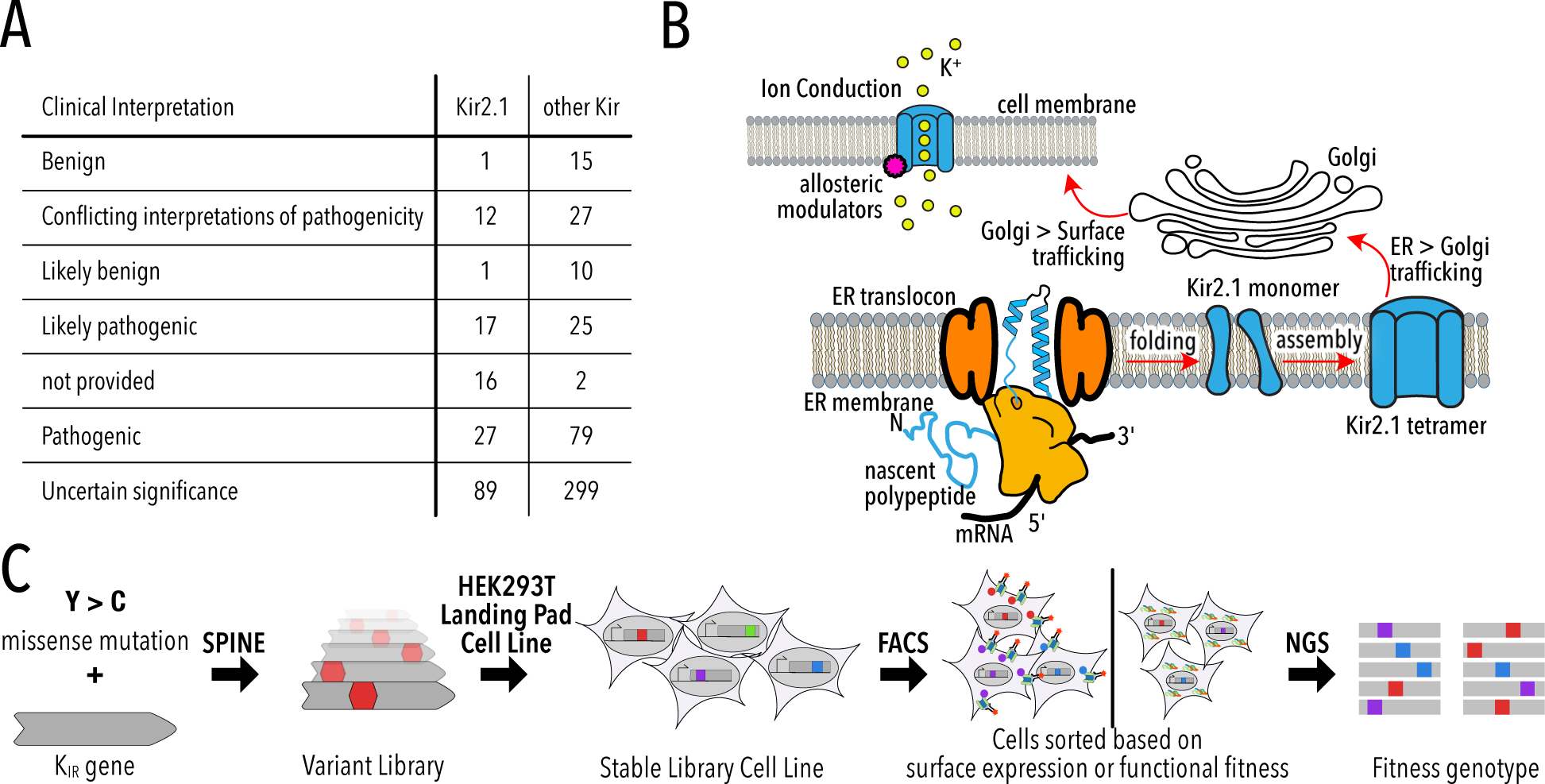
Deep Mutational Scanning to improve clinical variant prediction. **(A)** Clinically observed variation in Kir2.1 and other Kir and their interpretation of pathogenicity. **(B)** Schematic summarizing the various processes that myst work correctly for Kir2.1 to fulfill its role in cellular physiology. **(C)** Schematic summarizing DMS and multiparametric assessment of phenotype. Mutations are introduced into Kir2.1 using SPINE. A stable single-copy insertion library is generated by BxBI-mediated recombination in HEK293T. Cells are sorted based on channel surface expression or function (K^+^ conductance) as determined by antibody labeling of an extracellular FLAG tag or voltage-sensitive dye, respectively. Genotypes of each sorted cell population are recovered by NextGen Sequencing (NGS).

Despite a central role for Kir trafficking and function in disease etiology, there are no large-scale studies that comprehensively study sequence determinants of KIR trafficking and function. Previous studies focused on a small subset of common natural variants, while systematic variation is needed for a global picture of Kir trafficking and function. Beyond explaining disease, large-scale mutational studies can reveal intrinsic properties of proteins ^32–34^, determine protein structures ^35^, and inform mechanistic models of protein structure function relationships ^36^.

Here, we combine programmable mutagenesis library generation with multi-parametric sequencing-based assays for surface expression and function of Kir2.1. Our data reveal distinct determinants for Kir trafficking, folding, tetramerization, and function. The phenotype of most clinically observed variants can be explained by impacting function, which may be related to the essential role of Kir2.1 in human physiology. We also find further support for the hypothesis that a hierarchical organization of ion channels structure balances stability and flexibility for folding and function ^34^.

### A multi-parametric deep mutational scan of kir2.1

For Kir2.1 to fulfill its roles in cellular physiology, it must be ER targeted, folded, tetramerized, and surface trafficked (**Fig. 1B**). Once trafficked to the cell surface, Kir2.1 must be sensitive to Phosphatidylinositol 4,5-bisphosphate (PIP_2_) ^1, 37^, undergo gating associated conformational transitions (pore opening and closing), and selectively conduct K^+^.

To probe how Kir2.1 mutations affect these processes, we use a programmable deep mutagenesis approach (SPINE ^38, 39^) and mutated residues 2-392 of mouse Kir2.1 (Uniprot P35561) to every other amino acid (**Fig. 1C**). We included synonymous mutations in 20 Kir2.1 positions as an internal standard to determine wildtype fitness. From this DMS library, we generated a stable single copy library cell line using BxBI-mediated recombination ^40^ for stringent genotype-phenotype linkage. Endogenous channel currents in HEK293 are negligible when compared to those of exogenous channel driven from a strong constitutive promoter (CAG) ^41^.

Since mutations can affect any of the abovementioned processes, multi-parametric assessment of phenotypes is required to develop a comprehensive mechanistic understanding of Kir2.1 variation. We focused on separating a mutations effect on secretion, folding, and trafficking (‘surface expression’) and allosteric regulation and ion conductance (‘function’). Considering the library size of variants (391x19 = 7,429 variants), assays that assess surface-expression and function must be high-throughput. Cell-based assays coupled to fluorescently activated cell sorting that can separate cell population (and the genotypes encoding in population) based on a phenotype reported by fluorescent reporters are immensely flexible.

Mutations that interfere with secretion, folding, or trafficking will cause reduced Kir2.1 surface expression ^42–44^. To measure Kir2.1 surface expression in cells we inserted an epitope tag into the extracellular loop (T115 ^45^). Cells with surface trafficked variants can be separated from poorly expressed variants based on antibody-labeling and fluorescence-activated cell sorting (FACS) (**Supplementary Fig. 2**). Phenotypes can be linked to genotype by NGS. We’ve extensively used this method in FACS-based genotype-phenotype assays ^34, 38, 46^.

**Figure 2.**
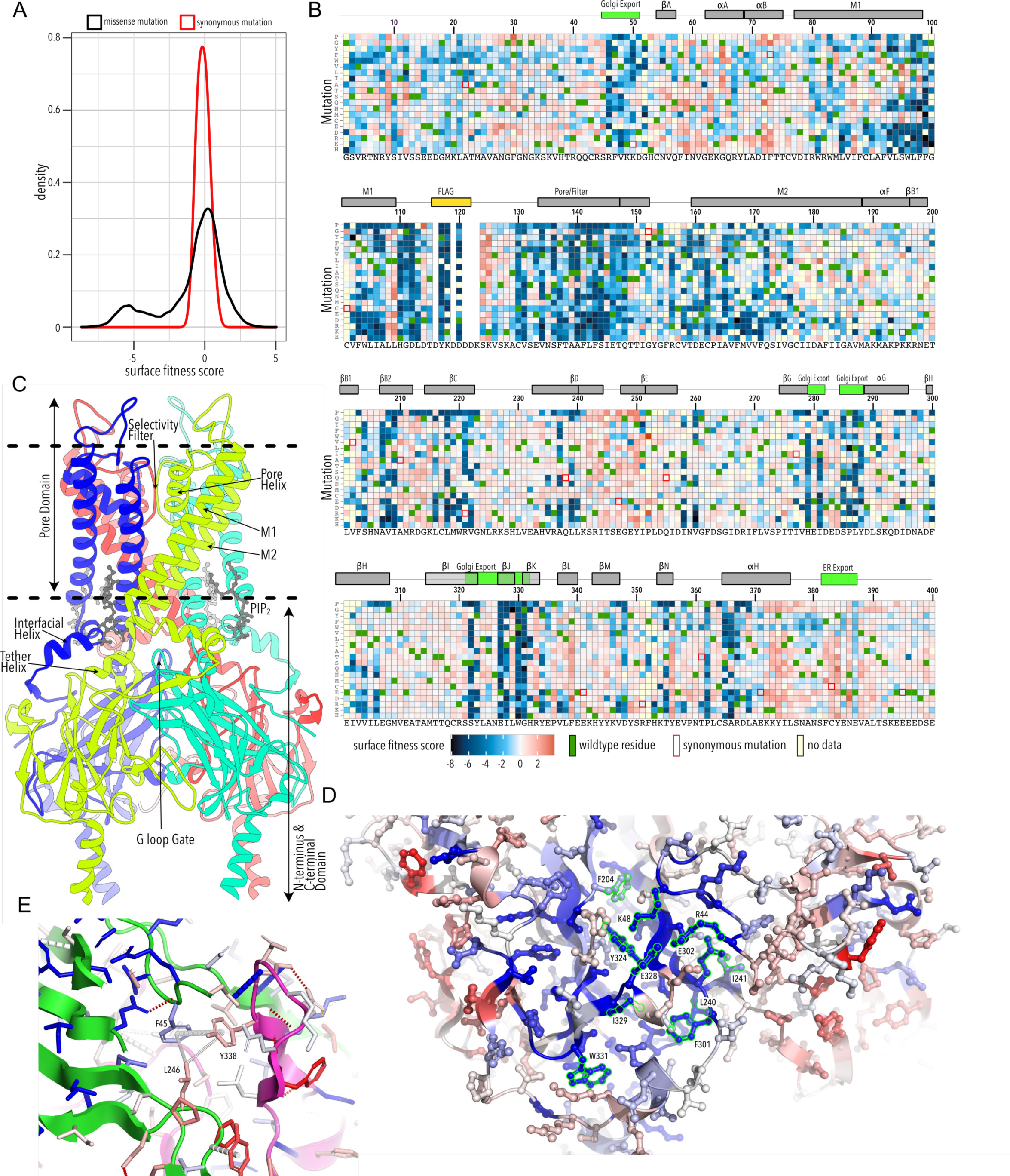
Surface Expression Fitness. **(A)** Distribution of surface expression fitness scores for synonymous mutations (red line) and missense mutations (black line). **(B)** Heatmap of surface expression fitness scores (gradient from blue to red). Wildtype residues are indicated by green boxes. Missing data is indicated by yellow boxes. Synonymous mutations are indicated by red box outline. **(C)** Cartoon of Kir2.2 (PDB 3SPI, 70% identity with Kir2.1). Monomer chains show in different colors. Lipid bilayer boundaries are indicated by dashed lines. **(D)** Close-up of core beta sheet of the CTD with surface expression fitness mapped onto each residue as a gradient from blue to red. Residues that comprise the Golgi export signal are outlined green. **(E)** Interface close-up of neighboring subunits (green and magenta cartoons). Sidechain are colored by surface fitness score. Disrupting carbon–π and π–π interactions interactions between F45, L246, and Y338 does not uniformly decrease fitness.

Functional Kir2.1 hyperpolarizes HEK293 cells by driving the resting membrane potential (RMP) towards the reversal potential of K^+^ (-80mV) while non-functional Kir2.1 variants do not affect the RMP (-35mV). We can measure RMP with voltage sensitive dyes, such as the FLIPR Blue, a concept that we and others demonstrated in optical Kir functional assays ^46, 47^. Changes in membrane voltage alter dye membrane partitioning, and therefore extinction coefficient and fluorescence. Hyperpolarized cells expressing functional Kir2.1 can be separated from more depolarized cells expressing non-functional Kir2.1 by FACS using FLIPR dye fluorescence (**Supplementary Fig. 3**). As with surface expression, phenotype can be linked to genotype by NGS.

**Figure 3.**
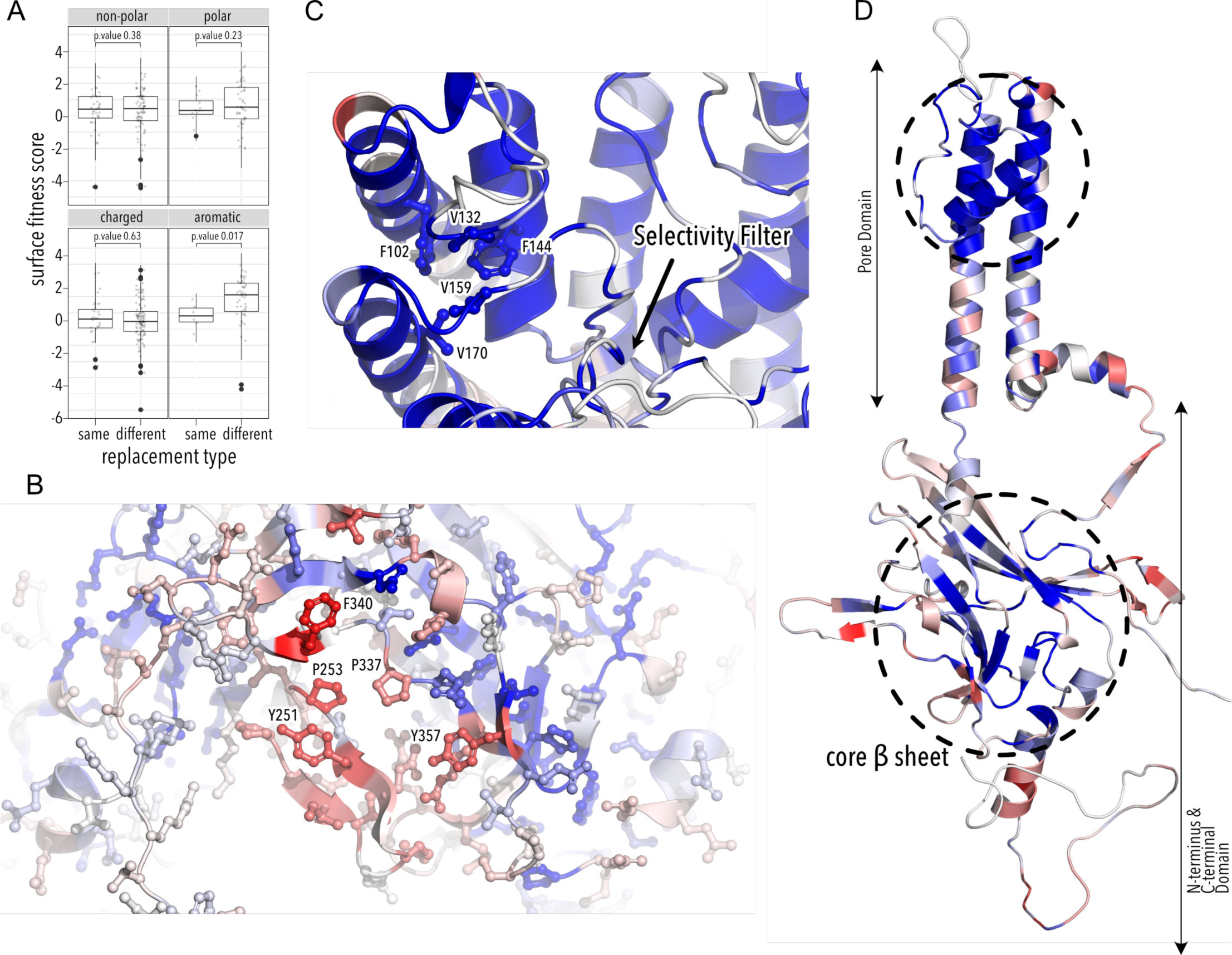
Determinant of improve surface expression and Kir2.1’s biogenic folding unit. **(A)** Boxplots showing surface fitness in the βDE and βLM loop if the wildtype residues of a specific type (e.g., glutamate, charged) is mutated to another residue of the same type (e.g., Aspartate) or a different type (e.g., Tryptophan). Median is marked with a thick line, the vertical length of the box represents the interquartile range (IQR), upper fence: 75th percentile +1.5 × IQR, lower fence: 25th percentile −1.5 × IQR, outlier points are shown as solid black circles. All data points are indicated by transparent dots. Only replacing aromatic residues significantly increase surface expression. Significance is tested using one-sided t-tests; p-values are indicated for each comparison. **(B)** Close-up of surface expression fitness scores (gradient blue to red) in the βDE and βLM loop. **(C)** Close-up of surface expression fitness scores in the pore domain. Residues that are part of the putative biogenic unit of Kir2.1 are show as sticks. **(D)** Surface expression fitness scores mapped onto a single Kir2.1 monomer highlighting the importance of the biogenic folding unit in the pore domain and the core beta sheet of the CTD.

### A global view of trafficking determinants

To learn how amino acids contribute to and mutations alter Kir2.1 trafficking and folding, we sorted and sequenced the Kir2.1 DMS library based on surface expression by FACS-seq. Surface trafficking fitness was determined for 6,898 Kir2.1 variants (93%). Biological replicates for subpools were highly correlated (Pearson correlations 0.85-0.94, **Supplementary Fig. 4**) and shows read depth is excellent (greater than 30-fold at most positions, **Supplementary Fig. 5**; **Supplementary Table 1A**). Surface expression fitness scores and standard errors were calculated using Enrich2 ^48^. Enrich2 calculates log enrichment by fitting a weighted regression based on changes in mutation frequency across samples within an experiment. Median fitness error relative to measurement rang was low (7.5%) in the surface expression assay based on synonymous mutation standard deviations (**Fig. 2A, Supplementary Fig. 6**). Surface fitness follows a bimodal distribution with one population of mutations centered at wildtype fitness and the other population strongly decreasing surface expression (**Fig. 2A**).

**Figure 4.**
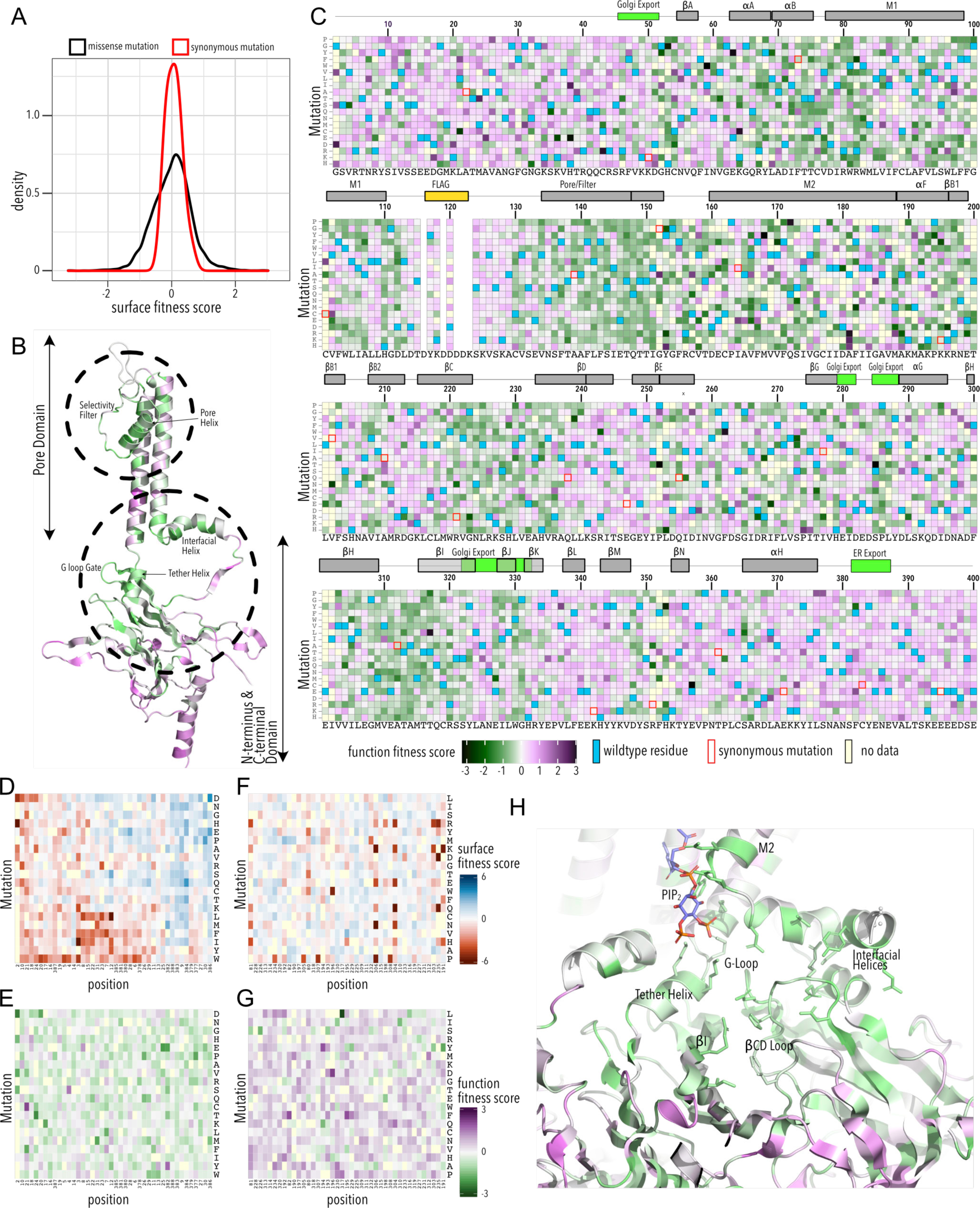
Function Fitness. **(A)** Distribution of function expression fitness scores for synonymous mutations (red line) and missense mutations (black line). **(B)** Function fitness scores mapped onto a single Kir2.1 monomer highlighting the importance of the pore helix and selectivity filter, and the structural elements are the TM/CTD interface (e.g., interfacial helix, tether helix, G-loop gate) for function. **(C)** Heatmap of function expression fitness scores (gradient from green to magenta). Wildtype residues are indicated by blue boxes. Missing data is indicated by yellow boxes. Synonymous mutations are indicated by red box outline. (**D-G)** Heatmap of fitness score in regions that contain bona-fide trafficking signal (**D,E;** for surface and function scores, respectively) or the PIP_2_ binding site (**F,G;** for surface and function scores, respectively). (**H**) Closeup of function fitness at the TM/CTD interface. Allosteric ligand PIP_2_ is shown as sticks.

**Figure 5.**
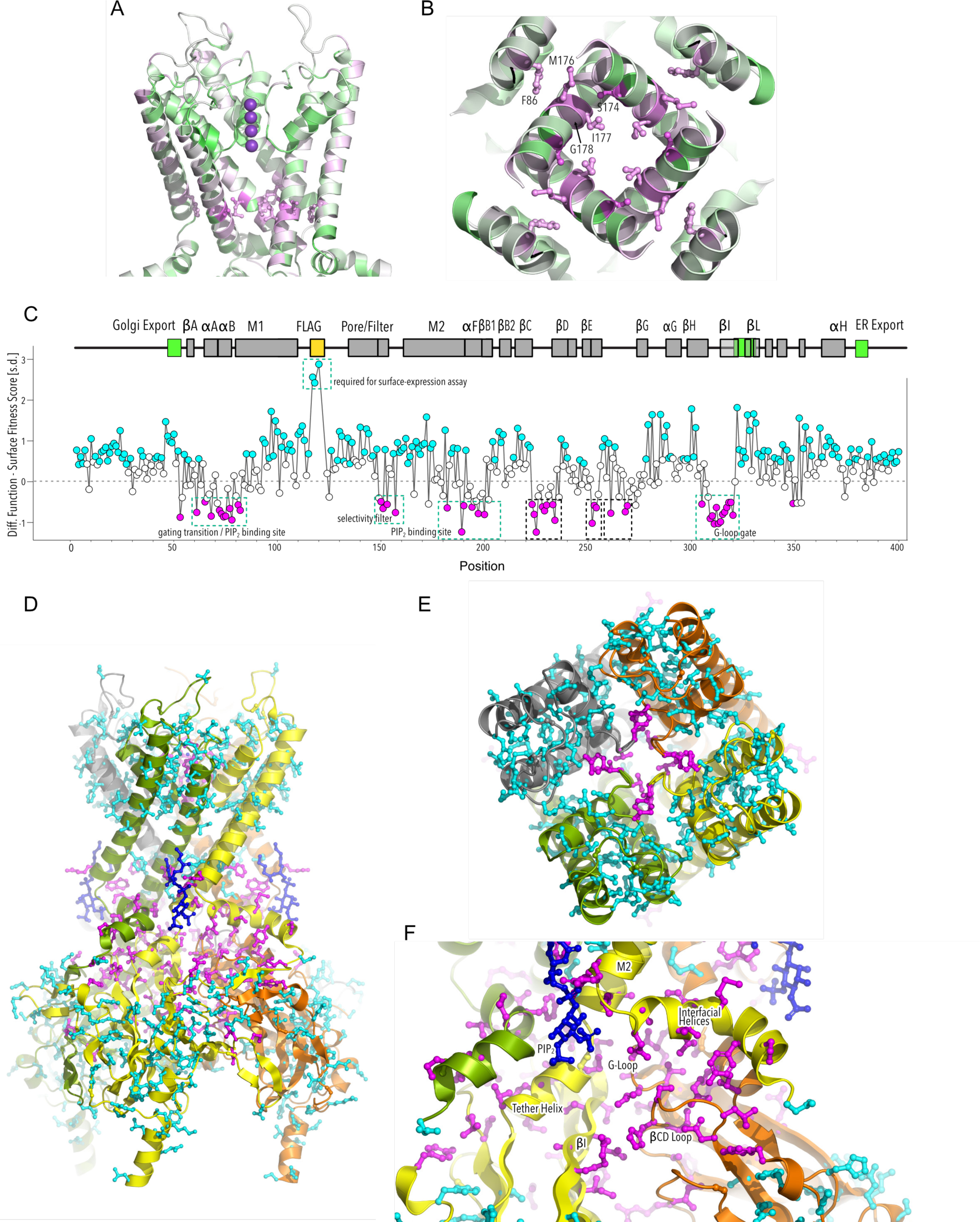
HBC gate mutations and regions with distinct roles stability and function. **(A)** Function Fitness (gradient from green to magenta) in the pore domain. One subunit is removed for clarity. K+ ions in the selectivity filter a show as purple spheres. Sidechain of residues that are enriched for activating mutations are shown as sticks. **(B)** Detailed view of the HBC gate. **(C)** Difference between z-score normalized mean Function and Surface Fitness scores mapped by position. Filled circles (cyan or magenta) indicated significant differences (two-sided moderated t test p-value < 0.05). Kir2.1 secondary structural element are shown as gray boxes; trafficking signals are colored green. **(D-F)** Residues with significant function/surface fitness score differences are mapped onto the Kir2.2 structure (PBD 3SPI). Backbone cartoons are colored by chain. Residues with mutation that predominantly reduced function are shown in magenta. Residues with mutation that predominantly reduced surface expression are shown in cyan. PIP_2_ is shown in blue. Overview **(D)**, close-ups of selectivity filter **(E)** and TM/CTD **(F)**.

**Figure 6.**
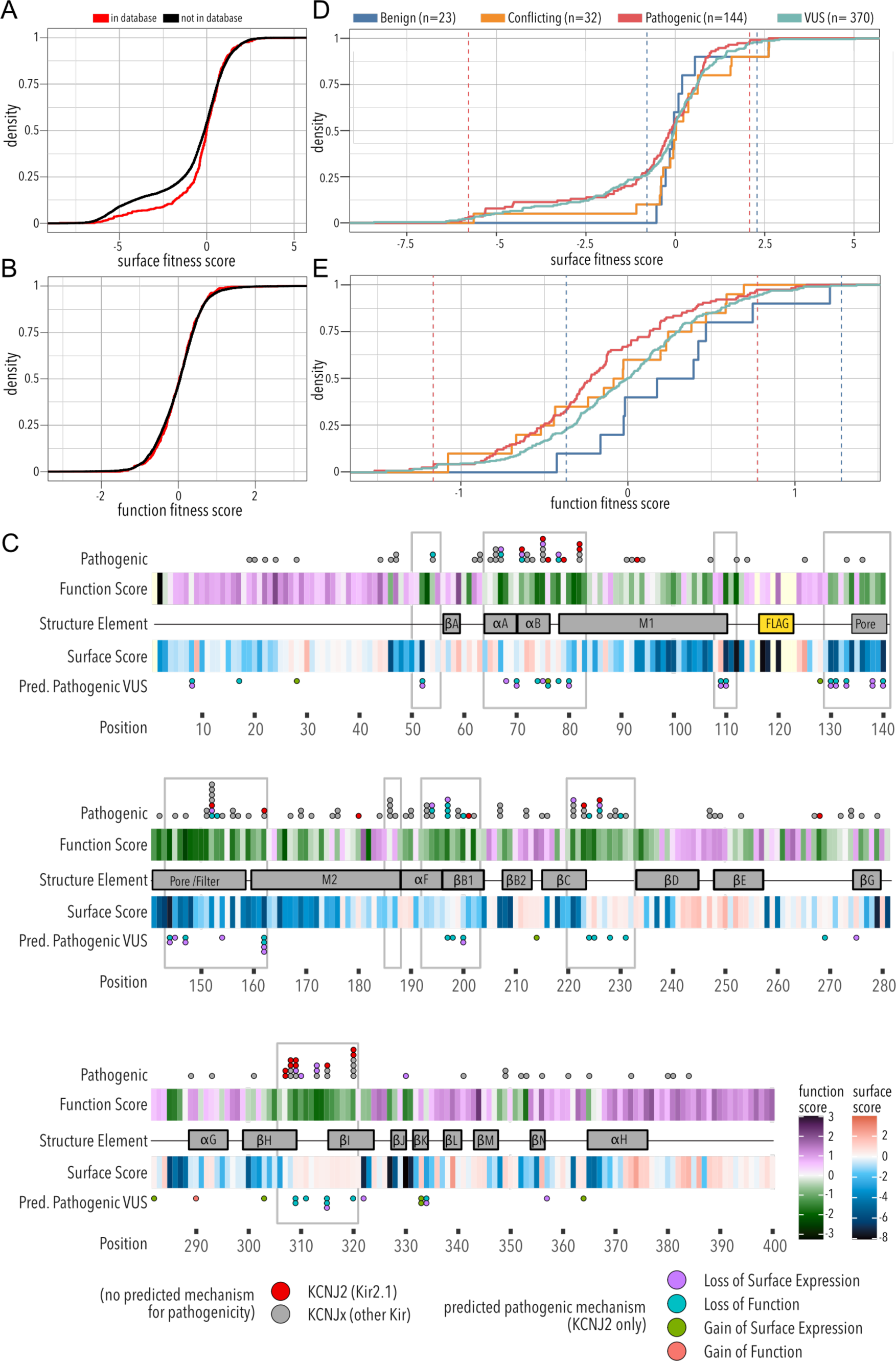
Folding and trafficking variants in clinical databases. Cumulative distributions of surface **(A)** or function **(B)** fitness scores for variants that are represented in Clinvar and dbSNP (red line) or that are not (black line). **(C)** Line plot by position of mean function or surface fitness scores (gradient green to magenta and blue to red, respectively). Kir2.1 secondary structural element are shown as gray boxes. Pathogenic variants are mapped as circles with fill color indicating predicted pathogenic mechanism. Mutation hotspots are indicated by gray outlines. Distribution of surface fitness **(D)** and function fitness **(E)** by ClinVar pathogenicity assignment. Vertical dashed lines represent 2.5^th^ and 97.5^th^ percentiles bounds of expert reviewed pathogenic variants (red) and benign (blue), respectively.

As expected, mutations are highly deleterious within the Flag-tag used for surface labeling and known trafficking signals (**Fig. 2B**). Substituting aromatic residues with charged residues strongly decreases surface expression within the ^81^WRWMLLLLF^88^ motif that mediate interaction with Caveolin-3 ^49^. Glutamates within the di-acidic ER export signal, ^382^FCYENE^38750^, are quite sensitive to mutations with the second glutamate being far more sensitive. Interestingly, negatively charged mutations, N386E or N386D, between the di-acidic motif increase surface expression.

A Golgi export signal comprised of two residues patches at Kir2.1’s N-and C-termini interacts with the AP1 adaptin complex to target Kir2.1 to clathrin-coated vesicles for Golgi to plasma membrane trafficking ^25^. Others have used alanine mutagenesis screens to identify sequence determinants for Golgi export. First, Ma et al. ^25^ defined the minimal signal motif comprised of hydrophobic residues (I321, W323, Y316) and a juxtaposed electrostatic interaction between R46 and E327. Later, Li et al. ^18^ included salt bridges between basic residues in the N-terminus (R44, R46, K50) and C-terminus (E3201, E327), and hydrophobic residues (F203, L239, I240, F300). This structured trafficking signal is formed when the CTD has folded correctly to include a hydrophobic cleft, an adjacent cluster of basic residues, and a network of interactions between termini and adjacent subunits. Ma et al. suggested this structured trafficking signal predisposes Golgi exit on correct protein conformation; our comprehensive mutagenesis suggests this quality control mechanism may extend beyond previously identified sequences. It appears to involve all core beta sheet in the CTD (βD1, βH, βI, βJ, βK, βB2, βC, βG) and helices αG and αF (**Fig. 2B,C**). Across CTD secondary structure elements, we find a distributed network of residues that, when mutated, result in reduced surface expression (**Fig. 2D**). Changes to the packing of the hydrophobic core of the CTD will likely impact the Golgi export motif orientation. We hypothesize that the Golgi export signal is a reporter for folding of the entire CTD. Several lines of evidence support this hypothesis: IgG-like domains are a beta sheet sandwich with conserved hydrophobic residues at sheet interfaces that are crucial for fold stability ^51, 52^, computational mutagenesis in Kir1.1(ROMK1) showed these residues are crucial for fold stability, and experimental mutagenesis found mutations strongly reduces inward rectifier surface expression ^15, 53^. Residues in this network belong to the same monomer; disrupting interactions between residues belonging to neighboring monomers (e.g., carbon–ð and ð–ð interactions between F45, L246, Y338; **Fig. 2B,E**) do not uniformly reduce surface expression. Although Kir are presumably already assembled into tetramers ^54^, our data suggests the conformational checkpoint for Golgi export requires properly folded monomers not correctly assembled tetramers.

Sites enriched for beneficial or deleterious substitutions could point to additional mechanisms for Kir2.1 trafficking control. Along those lines, we observe that substitution of N-and C terminal residues (G2–G31, S377-S381) with polar mutations have neutral trafficking phenotypes, while aromatic or hydrophobic mutations tend to decrease surface expression. While we do not know the underlying mechanism, our data suggests disordered N-and C-termini are important in targeting Kir2.1 to the cell surface.

Mutations along βB2(M211–K215), βDE(S242–P252), βK(R333, P336), βL(F339), βM(Y345), βN(Y366), and αF helix(K372–L376) tend to improve surface-trafficking above WT especially when aromatic residues are replaced with any non-aromatic amino acids (**Fig. 3A**, t test p-value 0.017). These sites form a patch on the outer face of the CTD domain where two subunits interdigitate (βDE from one subunit, the remaining elements from another subunit; **Fig. 3B**). In this region beneficial mutations have no clear physicochemical preferences beyond non-aromatics. While no known trafficking signal matches, aromatic residues are often binding interfaces ^49, 55^. Perhaps mutations alter trafficking patterns by disrupting protein interactions (e.g., reduce ER retention, reduce forwarding to lysosomal recycling).

The agreement with earlier trafficking studies in Kir2.1 shows that our high-throughput surface expression assay works. This allows us to establish a global view of sequence/trafficking relationships in Kir2.1. Comprehensive mutagenesis allows us to validate and fill in the gaps on existing models of channel trafficking ^18, 25^ while discovering new potential trafficking determinants.

### A cotranslationally-folded biogenic unit in Kir2.1

By examining structured regions sensitive to mutations, we identify determinants of Kir2.1 folding. In the pore domain, we find the lower halves of M1 and M2 are surprisingly tolerant to mutation, in particular residues whose sidechains project towards the pore cavity. At the base of the pore domain, mutations are highly deleterious within a cluster of hydrophobic residues (W81, L85, I178, F182, I187, M191) that form inter-monomer interactions between M1 and M2 helices. The top of the pore domain –a region comprised the upper halves of M1, M2, pore helix, and selectivity filter– is extremely intolerant to substitutions. This includes the M2-pore helix loop (e.g., H110–E133), whose amino acid sidechains contribute to tight packing centered around a hydrophobic pocket formed by F103 (M1), F143 (pore helix), V169 (M2), and V131 M1-pore helix loop (**Fig. 3C**). The same region contains two highly conserved cysteines (C130, C162) that form a disulfide bond which is critical for folding, not function (K^+^ conductance) ^56–58^. Others found P156, at M2’s apex, is essential for efficient surface trafficking in Kir2.2 ^59^. In agreement with folding critical roles, all substitutions in these positions are deleterious to surface expression.

Glycines in the selectivity filter (^147^TQTTIGYG^154^ in Kir2.1; conserved residues are underlined), which are absolutely conserved in all K^+^ channels, are immutable. Because glycine is achiral, it is the only canonical amino acid that can satisfy the unusual conformation of the selectivity filter main chain, which alternates between left-handed and right-handed α-helical dihedral angles ^60^. Substitutions of either T150 (which occupies unfavorable backbone torsion angle conformation in the Kir2.2. crystal structure ^37, 56^) or Y153 (which forms π–π interaction to a neighboring subunit) in the selectivity filter with glycine significantly increase surface expression. This suggests the peculiar conformational requirements of the selectivity filter are a constraint in channel biogenesis.

Based on the extreme mutational sensitivity, high contact density of side chain packing in the Kir2.2 crystal structure, and known disulfide-mediated fold stabilization, we propose that the top halves of M1 & M2, pore helix, and the M2-pore helix loop promote tertiary structure formation within nascent Kir2.1 monomers (perhaps in the ER translocon) to establish correct topology and promote membrane insertion. This may be a general feature of K^+^ channel folding, as a similar “cotranslationally folded biogenic unit” was proposed for Kv1.3, in which S5/pore loop/S6 interact to establish a “reentrant pore architecture” and correct topology early in channel biogenesis without requiring tetramerization ^61, 62^. Interestingly, residues from neighboring subunits interact with this putative biogenic unit in the fully assembled channel (e.g., selectivity filter residues Y153, which forms a ð–ð interaction with F143 and F155, carbon–ð interaction with E133) are relatively tolerant to mutations. This supports the idea that early trafficking-critical folding events involve only intra-monomer interactions.

### Regions involved in gating transitions to do not contribute to fold stability

From the perspective of surface-trafficking as the measured phenotype, several regions of Kir2.1 are tolerant to mutations. This includes the interface between pore and CTD (comprised of slide helix αA/αB, inner helix gate (I183–M188), and tether helix αF), the G-loop (βH-βI loop), and the residues lining the CTD cavity (loops between βC-βD and βE-βG) at the interface between different subunits (**Fig. 3D**). Interestingly, these are involved in gating associated conformational changes ^37, 63^ induced by the binding of PIP_2_, Kir2.1 allosteric activator (two-sided Fisher’s Exact Test, p-value 8.091e-05, odds ratio 2.47). An explanation for high mutational tolerance in gating structural elements is that they have higher conformational flexibility required for interconversion between different gating states. They are therefore unlikely to contain motifs recognized by folding-based quality control mechanisms.

Kir2.1 appears to have an internal organization into contiguous regions involved in fold-stability (biogenic folding unit, IgG-like CTD) and gating transitions (TM/CTD interface, CTD subunit interfaces) with distinct mutational tolerance. To understand the relationship between folding and function, we assayed K^+^ conductance as a second phenotype for the DMS library.

### Functional phenotype assays map molecular determinants of conduction and PIP_2_ sensitization

To assay variant function, we sorted the Kir2.1 DMS library (in duplicate, on separate days) into based on voltage-sensitive dye fluorescence, sequenced, and calculated fitness using the same Enrich2 package. This assay is based on premise that Kir2.1 decreases RMP resulting in decreased fluorescence of the FLIPR dye. Functional fitness was determined for 6,944 Kir2.1 variants (93.4%), with similar replicates (Pearson correlations 0.86-0.93, **Supplementary Fig. 7**) and excellent read depth (greater than 30-fold at most positions, **Supplementary Fig. 8**; **Supplementary Table 1B**). Based on synonymous mutation standard deviations (**Fig. 4A**), median fitness error was low (15.3%). Unlike the bimodal distribution for surface-expression fitness, functional fitness is unimodal, with most mutations neutral (**Fig. 4A**); there is no distinct low function population likely due to lower dynamic range of the voltage sensitive dye compared to fluorescent antibody labelling (**Supplementary Figs. 2-3**). Function fitness is the combination of surface expression (number of channels on surface) and individual channel functional properties (open probability, single channel conductance, K^+^ selectivity, etc.). Furthermore, as our assay is measuring steady-state RMP and even partially functional Kir2.1 will eventually hyperpolarize the cell. This means we must interpret functional fitness in context of surface expression fitness.

Many variants in the pore domain have low functional fitness scores (e.g., pore helix, selectivity filter, **Fig. 4B**). For example, Y153 mutations, which have relatively tolerant surface fitness, strongly reduces function (**Fig. 4C**). Y153 interacts through π-π interactions with F143 and F155 on a neighboring subunit and mutations may preclude correct selectivity filter geometry to conduct K^+^ ions. The conservative substitution Y153F has a neutral phenotype, in agreement with earlier electrophysiology studies ^64, 65^. Many positions along the pore helix that form a hydrogen bond network to support a conductive and K^+^ selective state ^66^ are very sensitive to mutation. Mutations of conserved hydrophobic residues in the CTD core beta sheet (e.g., W220) and the Golgi export signal (e.g., hydrophobic cleft residues Y316, W323, and salt bridge R45, E327) are deleterious as predicted by their effect on surface-expression.

Disordered N and C termini including the ER export signal FCYENE had moderately low surface-expression fitness but their functional fitness is mostly near neutral (**Fig. 4D,F**). This suggests mutations within the disordered regions only impact trafficking of otherwise properly folded and functional channel. Mutations within the unstructured termini they are deleterious to surface-expression and but neutral to individual channel function.

PIP_2_ binding at the TM/CTD interface induces a disorder-to-order transition in the tether helix, contracting it and thereby translating the CTD toward the pore domain. The engagement between and CTD and pore domain allows the G-loop to wedge into the pore domain and the inner helix activation gate to open ^37^. PIP_2_’s binding site includes pore domain residues (R80, W81, R82, K190) and CTD residues (K186/193, K188/195, K189/196, R65, R219/226, R197). As expected, surface-expression fitness was neutral and functional fitness was negative (**Fig. 4E,G**). By focusing on mutation-sensitive positions, we can trace secondary structural elements that couple the PIP_2_ binding site to the Kir2.1’s gate, the G-loop ^67^. We find residues that directly interact with PIP_2_ in the pore domain (R80, W81, R82) and tether helix (K190, K193, K195, K196). We find additional residues in the tether helix (R197), βB1 (L202), and interfacial helix (R65) interact with the allosteric coupling βC-βD regions. Residues are highly sensitive in the entire βC-βD and its interactions including a hydrophobic interface between βC and βI with each subunit, several interactions couple to the βCD loop (e.g., H229) and βEG loop (e.g., F270) of one subunit to the βH (I305), βI (R320), and the G-loop gate of another subunit (**Fig. 4H**).

Taken together, deep mutational scanning reinforces a central role of the βCD loop propagating PIP_2_ binding at the TM/CTD interface to the G-loop gate. Molecular dynamics simulations uncovered an extended network of interactions between N-terminus, tether helix, CD loop keep the G-loop gate in a closed state ^63^. Electrophysiology demonstrated that mutations in R219, which is conserved in the KIR family, attenuate PIP_2_ affinity ^68^ and causes Anderson Tawill Syndrome ^69^. Mutating the corresponding residues in GIRK2/Kir3.2 (R201A) constitutively activates channels, presumably by forcing the βCD loop into conformation that mimics the G-protein activated state ^70^.

In GIRK2, an aspartic acid in βCD (D228) is a key residue in mediating regulation by intracellular sodium (an important physiological role of GIRK2). This Na^+^ binding site propagates conformational changes in the CTD’s β strands after Gβγ binds to βDE and βLM loops, displacing the βLM loop upward to interact with the N-terminus and βCD loop, ultimately stabilizing the open conformational of the G loop gate ^71, 72^. The corresponding wildtype amino acid in Kir2.1, which is not regulated by Na^+^, is N224 (**Fig. 4B**). Interestingly, substitution with aspartate increases Kir2.1 function, perhaps by recapitulating the Na^+^ binding site, which also involves coordination from main chain carbonyls (R226, K227, S228). The function increasing effect of N224K (and the deleterious effect of N224R) could be explained through the Na^+^-independent stabilization of the βCD loop through hydrogen bonding between the protonated epsilon-amino group of lysine (mimicking Na^+^) and the abovementioned carbonyl oxygens.

Another striking feature revealed by functional assays are function-increasing mutations near the helix bundle crossing gate of Kir2.1 (**Fig. 5A**). Previous studies suggested a kink occurring at G176 in TM2 widens the pore at the helix bundle crossing (HBC) allowing for K^+^ to pass ^73–75^. As expected, mutating G176 decreased function fitness. Substitution in residues below the kink-inducing G176 and above the actual bundle crossing (V187), increases functional fitness. This effect is strongest in A173, I175, I176, G177 (corresponding to S174, M176, I177, G178 in Kir2.2; **Fig. 5B**). Together, residues form a cuff above the HBC gate coupling TM2 from different subunits. The cuff may stabilize the unkinked conformation of TM2 to decrease open probability. Consistent with this idea, mutating cuff residues to large charged amino acids has the greatest function­increasing effect (**Fig. 4C**). Perhaps mutating to large, charged residues disrupts nearby hydrophobic interactions networks that stabilize the HBC close state. The HBC gate is also stabilized by hydrophobic contacts to the TM1 of an adjacent subunit ^76, 77^. Consistent with TM1 interactions forming a gating energy barrier, mutating interacting TM1 residues (F88; F86 in Kir2.2) also increases function fitness.

To identify regions that are important for function and not surface expression, we z-scored and subtracted surface expression from function fitness. We used t-tests with pooled variance estimated from all mutations to test significance of difference in fitness and function scores for each position (**Fig. 5C**). For many positions, surface scores were significantly lower than function scores (**Fig. 5C, cyan circles**). This subset includes the FLAG tag that is required for surface expression assay, but also Golgi export motifs, transmembrane helices, and CTD core beta sheets. For another subset of positions, function scores were significantly lower than surface-expression scores (**Fig. 5C**, magenta circles). This includes sites involved in gating (PIP_2_ binding, G-loop gate), and K^+^ conduction (selectivity filter). By making a binary assignment (based on p-value) whether a position belongs to one subset or the other and mapping this assignment onto the Kir2.2 structure, we again find structural organization around elements that are important for fold stability (TM, CTD beta sheets) and function (PIP_2_ sensitization, gating, K^+^ conduction) (**Fig. 5D**). This distinct organization is clearest in the pore domain, where all interactions within a subunit are driving surface expression score (**Fig. 5E**, cyan sticks), while residues mediating subunit interactions (which is required for stability of the selectivity filter) are driving function score (**Fig. 5E**, magenta sticks). Other residues that drive function scores are distributed along an interaction network that connects the PIP_2_ binding site (M2, αF) with the slide helix (αA/B), βCD loop, βEG loop, and the G-loop gate (βHI loop) (**Fig. 5F**). Intriguingly, this network involves interfaces between adjacent subunits (N-terminus<>βLM loop interface, βI<>βCD loop, βD<>βEG loop). Subunit interactions determining gating is consistent with recent crystal structures of a forced-opened Kir2.2 in which inner helix gate opening requires CTD subunits to move relative to each other ^77^.

Overall, the functional assay allows us to validate and test existing models of Kir2.1 gating while also providing further evidence that inter-subunit interactions serve as an energy barrier to bias Kir2.1’s conformational ensemble towards closed states.

### Most pathogenic inward rectifier mutations are clustered within functionally important residues

A central goal in molecular genetics is identifying the mechanistic basis by which mutations cause disease. Multiparametric DMS studies that interrogate multiple phenotypes associated with variants are a promising strategy to answer this question. Based on the premise that most mutation affect protein stability, approaches such as Vamp-Seq ^78^ were developed as generalizable measures of protein abundance. In cases such as Kir2.1, where proper trafficking, localization, and gating are crucial for producing functioning proteins, measuring abundance likely won’t work. It is necessary to differentiate between mutations’ effects on folding, trafficking, and gating to learn the mechanism for how genotype affects Kir2.1 phenotype. To learn potential pathogenicity and mechanism of action for reported disease-associated mutations, we compared the surface expression and fitness data to clinically observed mutations.

Since the overall domain architecture and structure/function relationships are conserved within the inward rectifier family, we reasoned that the mechanisms underlying variant effects are conserved, as well. We therefore gathered missense variant effects reported in Clinvar and dbSNP for inward rectifiers. By aligning all human Kir, we assigned the corresponding Kir2.1 position to each variant (total variant count: 613) and noted whether wildtype amino acid matches between Kir2.1 and the aligned Kir (**Supplemental Table 2**). To test if variants in Clinvar or dbSNP is related to surface-trafficking or function, we compared the empirical cumulative distribution functions of trafficking and function scores variants in databases vs. those that are not. They differed for trafficking scores but not for function scores (two-sided Kolmogorov-Smirnov test p-values 4.28x10^-8^ and 0.51, respectively; **Fig. 6A**). This means being listed in genetic variation databases is related to Kir2.1 function and less related to trafficking. Variants with low surface scores in our DMS, are underrepresented in databases. A likely explanation is that Kir2.1 is essential for normal physiology and therefore under strong selection. Indeed, Kir2.1 homozygous knockout mice die 8 to 12 hours after birth ^79^. Kir2.1’s missense constraint score, based on exome and whole-genomes sequencing deposited in gnomAD ^80^, is 0.5 (i.e., 121 observed vs. 240 expected missense mutations based on background mutation rates). Variants that misregulate surface trafficking likely are more deleterious (e.g., abolishing all K^+^ conductance) than variants affecting function (e.g., gating kinetics), which is why they are extremely rare in the living human population. Consistent with this idea, exome and targeted sequencing in products of conception showed a significant enrichment of pathogenic variants associated with cardiac channelopathies in stillbirths ^81^. Furthermore, only one variant (W232C) of the 5 most surface-trafficking impairing positions on Kir2.1’s Golgi export motif (R44, S315, E320, W323, G333) is reported in Clinvar.

The correspondence between variant listing and Inward Rectifier function is also apparent when we annotate mean surface-trafficking and function scores with pathogenic variants (**Fig. 6C**). Across the board, hotspots enriched for pathogenic disease-associated mutation have low function scores, whereas their surface-expression scores are more varied. Variants of uncertain significance followed this trend. In addition to regions involved in PIP_2_-binding (αA/B slide helix, lower part of M1, αF), and propagating conformational changes to the G-loop gate (βB1, βCD, βHI loop), pore helix and selectivity filter are also hotspots for pathogenic variants. A notable exception to down-trending function in mutational hotspots is a cluster of disease-associated variants near the intracellular end of M2 (D180-A186). Interestingly, many Clinvar-reported variants (e.g., I183L) are associated with Short QT syndrome, a disease that can be caused by gain of function mutations in KNCJ2(Kir2.1) ^82^. Consistent with this clinical phenotype, many mutations increase function (compared to wildtype Kir2.1). Based on known structure/function relationship in inward rectifiers, we speculate that substitutions increase function by increasing open probability of Kir2.1 by disrupting the inner helix gate (I183–M188).

We further divided fitness scores by assigned clinical significance to estimate the fitness score bounds of expert-reviewed benign and pathogenic variants (**Fig. 6D,E**). For surface fitness, we find that >97.5% of known benign variant have a fitness score > -0.43. With this lower bound assumption for benign variants, we predict that any ‘variant of unknown significance’ (VUS) scoring lower will be pathogenic with a loss of surface expression phenotype. Conversely, the upper surface fitness score bound (10^th^ percentile) for expert reviewed benign was 0.95, and VUS with greater surface scores are predicted to have a gain of surface expression pathogenic phenotype. We applied the same reasoning to function fitness scores. Using this estimate, we can assign a predicted pathogenic phenotype (loss/gain of surface expression/function) to 69 of 370 (18%) Kir2.1 VUS **(Supplemental Table 2)**. The primary sequence location of these predicted pathogenic variants is shown in **Fig. 6C**. Many predicted loss of function variants localize to the slide & tether helices, G-loop gate, and selectivity filter/pore helix. Gain of surface expression variants map to the βBC and βK loop near the interface to βDE of a neighboring subunit. The putative biogenic folding unit (top of M1 & M2, pore helix) contains several predicted loss of surface expression variants. Interestingly, no expert-reviewed pathogenic variants map to this region, which suggests that DMS-based prediction can identify new mutation hotspots. We can also use estimated bounds of benign variants to predict the mechanism through which 27 out of 144 (18.8%) expert-reviewed pathogenic variants cause Kir2.1 disorders. Of those, eight pathogenic variants have a predicted loss of surface expression while 19 are predicted loss of function. In line with function-impairing variant being overrepresented in ClinVar, the odds ratio between predicted loss of function phenotype of known pathogenic variants vs. all predicted pathogenic variants is significantly different (two-sided Fisher’s exact test, p-value = 0.003144).

Taken together, we find variants with deleterious surface-score are underreported in available database, which we propose is due to the essential nature of Kir2.1 in human physiology. Using surface and function scores for expert-assigned variant, we can estimate bounds for benign variants, which then allows us to predict VUS that are above or below these bound to the be gain or loss of surface expression/function, respectively. We can also predict the mechanism of action through which known pathogenic mutations cause Kir2.1 disorders. This suggests that DMS-derived surface-trafficking and function scores can be useful to predict pathogenicity and underlying mechanism of inherited and de-novo mutations. Because of the conserved architecture of the Inward Rectifier family, assignment of mutation hotspots may extend to the entire inward rectifier family. Further corresponding DMS studies in other members of the inward rectifier family are required to separate generalize themes from idiosyncrasies. For example, Kir6.2 requires binding to the ABC-transporter SUR1 to be expressed to the cell surface and function. This means that assay-reported phenotypes for a subset of variation at the Kir6.2/SUR1 interface (involving αA/B slide helix, M1, βLM sheets) have greater bearing in Kir6.2-linked diseases (e.g., diabetes mellitus). The methods described here are a blueprint for these studies.

## Discussion

Deep Mutational Scanning (DMS) quantitatively links protein phenotypes to the genetic variation within the protein’s coding sequence. This style of systematic amino acid substitution can reveal protein function ^83–85^, determine protein structure ^86^, and explain protein behavior in healthy and diseased cellular contexts ^87^. Because protein don’t exist in isolation –within the context of a cell folding, trafficking, and function states can all be modified through protein interaction (e.g., chaperones) or ligand binding (e.g., allosteric modulators)– the phenotypic outcome of genetic variation can be multi-facetted. To start to explore the numerous impacts of mutations, we measure two phenotypes of Kir2.1 variation: surface-trafficking K^+^ conduction. Both assays are based on cell sorting mediated by fluorescent signals (antibody binding to an extracellular loop and voltage sensor dye). By measuring how variants are enriched or depleted, we assign quantitative fitness scores. Many other assay types are compatible with this approach, providing opportunities for even richer phenotypic description of ion channel variation. For example, a recently described approach, HiLITR ^88^, could provide more granular resolution about trafficking motifs and Kir2.1 localization in living cells. Spontaneous spiking HEK cells ^89, 90^, which co-express Kir2.1, a voltage-dependent Na^+^ channel (Nav1.7), the channelrhodopsin CheRiff, and genetically encoded voltage indicator QuasAr2, could be adapted to evaluate how ion channel expression levels and gating properties impact excitability ^91^. When assays are conducted with chemical screening, they may aid in the discovery of novel allosteric modulators, state-dependent blockers, and molecular chaperones to precisely treat channelopathies based on genotype. Integration of disparate assays into a common framework of protein variation and its role in structure and function will be challenging but efforts to standardize reporting and unified statistical frameworks for interpretation (e.g., Enrich2^48^, the Atlas of Variant Effects ^92^) are on the horizon.

Many of our findings with respect to mutational sensitivity of trafficking motifs, core structural elements, and regions of Kir2.1 involved in gating align with existing knowledge and biophysical intuition. Unlike previous more limited screens and intuition, our data is quantitative and enables data-driven approaches to construct global biophysical models of Kir2.1 structure and function. For example, the Golgi export signal comprised of two patches at the N-terminus and CTD was first described as a minimal set of hydrophobic residues along a CTD cleft and juxtaposed basic residues ^25^ and later expanded to adjacent sites ^18^. Our comprehensive screen shows this trafficking motifs may indirectly probe the correct folding of the entire β-sheet core of the CTD. It reveals scope of what this trafficking motif is reporting to the cellular quality control machinery; not simply the proximity of a few basic and hydrophobic residues, but proper conformation of the entire CTD and assembly into tetramers.

We observed contiguous regions with neutral expression fitness involved in gating transitions while regions involved in fold-stability were highly sensitive to mutations. Looking at the same regions in functional data, we find the inverse. This suggests two things. First, comprehensive assessment of Kir2.1’s phenotypes after perturbation (i.e., mutation) is a high-throughput method to annotate protein sequences into classes linked to specific functions (e.g., putative trafficking signals, folding units, etc.). Conceptually this is similar to other high-throughput biochemical approaches that probe sequence-function relationships, such as circular permutation profiling ^93^ or high-throughput enzyme variant kinetics measurements ^94^. Second, combining multiple assayed parameters is key to discover general organizational principles in Kir2.1, specifically two distinct structural regions with distinct roles in providing fold stability or dynamics required for gating transitions. Expanding to more measured phenotypes and integrating datasets may be the blueprint for ‘sequencing-based’ biophysics that probes protein function, folding, and dynamics through steady-state biochemical experiments.

Our DMS studies provide additional context for the mechanistic basis of structure/function relation (e.g., reaffirming the βCD loop’s role in propagating PIP_2_ binding at the TM/CTD interface to the G-loop gate). We find a biogenic folding unit that represents an early quality control step of reentrant pore architecture and correct topology likely while the monomer is in the translocon. A similar biogenic unit was described in voltage-dependent K^+^ channels, where it is stabilized by an extensive network of interactions ^61^. Furthermore, many ion channels have structurally homologous reentrant pore loop architectures and earlier studies suggested that the presence of an ‘aromatic cuff’ is a general feature of their biogenesis ^61, 95^. DMS studies may provide a path to probe the energetics and biophysics of stabilizing interactions and to test the hypothesis that hydrophobicity is a general stabilizing factor across reentrant pore loop architectures.

DMS may also shed light on how subunit interactions determine inward rectifier gating properties. We find mutations near the interface of subunits strongly increase Kir2.1 function. This is consistent with subunits roles in setting channel gating properties. Single channel patch electrophysiology of Kir2.1/2.2 heterotetramers showed that addition of a Kir2.2 monomer increases single channel conductance and decreases the open dwell time (τopen) ^96^. Perhaps this feature contributes to the observed differences in gating between inward rectifier homo-and heterotetramer, which differ in number and nature of subunit interactions.

Our finding that variants with deleterious surface-score are underreported in available database supports the emerging theme that many disease-causing variants are linked to trafficking defects ^13–16, 16–23^. Other large scale mutational analysis, as undertaken for the voltage-dependent K^+^ channel Kv11.1^97, 98^, has similarly shown that 88% of Long QT-linked variants have trafficking­deficient mechanisms. They also demonstrate that data-driven approaches outperform smaller, more limited studies that predicted normal trafficking for most mutants ^99^. Underrepresentation in variant databases has implications for clinical practice, since standards for the interpretation of sequence variants are heavily focused on null variants (nonsense, frameshift), population frequency (frequent variants are likely benign), predicted functional effect (is the variant in an important domain), and case evidence ^100^. In genes under strong purifying selection, such as KCNJ2/Kir2.1, missense mutations cause severe developmental defects (craniofacial structures, limb development ^79, 101^) likely contribute to or cause spontaneous abortions and pregnancy loss. These variants therefore never enter population-or clinical-associated variant databases. DMS studies are not limited by organismal viability to probe the structural and functional consequences of genetic variation. DMS could be useful to identify drivers of spontaneous abortions and the underlying mechanisms of recurrent pregnancy loss. By comparing our experimentally determined fitness scores to expert-reviewed assignment of variants effects, which are often based on smaller scale direct biochemical studies, we can estimate the bounds of fitness scores for benign and pathogenic variants. This allows us to make predictions about pathogenicity and mechanisms of action for variants of unknown significance (VUS) and known pathogenic variants, closing the loop between high-throughput assays, biophysical mechanisms underlying fitness scores, and clinical interpretation of human variation in ion channel genes.

These examples demonstrate that deep mutational scanning of Kir2.1 combined with two simple readouts, provides detailed insight into the mechanistic basis of ion channel folding, function under steady state conditions, and how these processes become misregulated in disease.

## Materials and Methods

### Kir2.1 DMS Library Generation

Into mouse Kir2.1 (Uniprot P35561), we introduced a FLAG tag into an extracellular loop (at position T115) and added a downstream expression marker miRFP670 co-expressed via a P2A sequence. For Kir2.1 residues 2-391, each wildtype amino acid was mutated to all other 19 amino acids weighted by their codon usage frequency in humans. In addition to these missense mutations, mutations synonymous to wildtype were included for 20 positions as a benchmark. We included missense mutation into the FLAG as a negative control. We generated this mouse Kir2.1 DMS library using the SPINE ^38^, which we briefly summarize as follows: SPINE based libraries employ synthesized pools of DNA oligos with mutations at each position of a gene. Due to high error-rates in oligo synthesis, the current maximum length for oligos is 230 base pairs. As most genes are longer than 230 base pairs, we break up our gene (8 section in the case of Kir2.1) and replace a subsection of the gene with a pool of mutated oligos. The oligos are designed with unique barcodes for amplifying out a specific subpools library, Golden Gate compatible BsmBI cut sites, and the mutation within a subsection of Kir2.1. OLS oligos were designed using the SPINE scripts on Github (https://github.com/schmidt-lab/spine). These scripts design oligo libraries, primers for amplifying oligo-sublibraries, and inverse PCR primers for adding compatible cut sites to the Kir2.1 plasmid.

All backbone were amplified using a 25 cycle PCR with GXL polymerase and 1 ng of backbone DNA as template. The PCR product was then gel purified. All oligo libraries were amplified using 25 cycles PCR with GXL polymerase and with 1 ul of the OLS library (resuspended in 1 ml TE) as template. To assemble the backbone and library DNA, BsmBI Golden Gate reactions were set up in 20 ul containing 100 ng of amplified backbone DNA, 20 ng of amplified oligo DNA, 0.2 μl BsaI-HFv2 (New England Biolabs), 0.4 μl T4 DNA ligase (New England Biolabs), 2 μl T4 DNA ligase buffer, and 2 μl 10 mg/ml BSA. These reactions were put in a thermocycler overnight using the following program: (1) 5 min at 42°C, (2) 10 min at 16°C, (3) repeat 40 times, (3) 42°C for 20 min, (4) 80°C for 10 min. This reaction was cleaned using a Zymo Research Clean and Concentrate 5 kit and eluted in 6 ul of elution buffer. The entirety of this reaction transformed in E. Cloni 10G electrocompentent cells (Lucigen) according to manufacturer’s instructions. Cells were grown overnight with shaking at 30°C to avoid overgrowth in 30 ml of LB with 40ug/ml kanamycin and library DNA was isolated using Zymo Zyppy miniprep kits. A small subset of transformed cells was plated varying dilutions to assess transformation efficiency and validate successful mutations. All libraries at this step yielded >300,000 colonies implying a 100x coverage (assuming 1/3 of the variants were perfect based on our previous analysis of Agilent OLS based libraries). Each sublibrary was combined at an equimolar ratio to make a complete library with all intended mutations included.

### Stable Cell Line Generation

To generate cell lines, we used a rapid single-copy mammalian cell line generation pipeline ^40^. Briefly, mutational libraries are cloned into a staging plasmid with BxBI­compatible *attB* recombination sites using BsmBI Golden Gate cloning. We amplify the staging plasmid backbone using inverse PCR and the library of interest with primers that add complementary BsmBI cut sites. Golden Gate cloning and subsequent transformation is conducted with BsmbI (NEB), T4 Ligase (NEB) following manufacturer’s using the same protocol as previously described for library generation. Completed library landing pad constructs are co­transfected (1:1) with a BxBI expression construct (pCAG-NLS-Bxb1) into (TetBxB1BFP-iCasp-Blast Clone 12 HEK293T cells) using Turbofect according to the manufacturer’s instructions in 6 wells of a 6-well dish. This cell line has a genetically integrated tetracycline induction cassette, followed by a BxBI recombination site, and split rapalog inducible dimerizable Casp-9. Cells were maintained in D10 (DMEM, 10% fetal bovine serum (FBS), 1% sodium pyruvate, and 1% penicillin/streptomycin). Two days after transfection, doxycycline (2 ug/ml, Sigma-Aldrich) is added to induce expression of our genes of interest (successful recombination) or the iCasp-9 selection system (no recombination). Successful recombination shifts the iCasp-9 out of frame, thus only cells that have undergone recombination survive, while those that haven’t will die from iCasp-9-induced apoptosis. One day after doxycycline induction, AP1903 (10 nM, MedChemExpress) is added to cause dimerization of Casp9 and selectively kill cells without successful recombination. One day after AP1903-Casp9 selection, media is changed back to D10+ Doxycycline (2 ug/ml, Sigma-Aldrich) for recovery. Two days after cells have recovered, cells are reseeded to enable normal cell growth. Once cells reach confluency, library cells are frozen in 50% FBS and 10% DMSO stocks in aliquots for assays.

### Surface Expression cell sorting

Thawed stocks of library cell lines were seeded into a 10 cm dish and media was swapped the following day to D10. Cells were grown and split before confluency to maintain cell health. Media was swapped to D10 + doxycycline (2 ug/ml, Sigma-Aldrich) two days prior to the experiment. Cells were detached with 1 ml Accutase (Sigma-Aldrich), spun down and washed three times with FACS buffer (2% FBS, 0.1% NaN3, 1X PBS), incubated for 1-hour rocking at 4°C with a BV421 anti-flag antibody (BD Bioscience), washed twice with FACS buffers, filtered with cell strainer 5 ml tubes (Falcon), covered with aluminum foil, and kept on ice for transfer to the flow cytometry core. Before sorting, 5% of cells were for processing and sequencing as a baseline control. Cells were sorted on a BD FACSAria II P69500132 cell sorter. miRFP670 fluorescence was excited with a 640 nm laser and recorded with a 670/30 nm bandpass filter and 505 nm long-pass filter. BV421 fluorescence was excited using a 405 nm laser. Cells were gated on forward scattering area and side scattering area to find whole cells, forward scattering width, and height to separate single cells, miRFP670 for cells that expressed variants without errors (our library generation results in single base pair deletions that will not have miRFP670 expression because deletions will shift the fluorescent protein out of frame ^38^), and label for surface expressed cells. The surface expression label gate boundaries were determined based on unlabeled cells from the same population because controls tend to have non-representative distributions. Example of the gating strategy for is depicted in **Supplemental Fig. 1**.

Cells were sorted based on surface expression into 4 populations (miRFP^high^/BV421^none^, miRFP^high^/BV421^low^, miRFP^high^/BV421^medium^, miRFP^high^/BV421^high^. We collected at least 2.1 million cells in each population to ensure a greater than 100x coverage for each sample in each subpool. The surface expression experiment was done in duplicate on separate days for two entirely independent replicates.

### Resting membrane potential cell sorting

Concurrently with preparing samples for surface labeling another sample of the same cells were prepared for sorting based on resting membrane potential. These cells were initially washed with FACS buffer and concentrated in this buffer for cell health prior to sorting. 30 minutes prior to sorting cells were resuspended in Tyrode (125mM NaCl, 2mM KCl, 3mM CaCl2, 1mM MgCl2, 10mM HEPES, 30mM glucose, pH 7.3) that contained FLIPR membrane potential dye with a Blue quencher.

Cells were also sorted based on resting membrane potential on a BD FACSAria II P69500132 cell sorter. miRFP670 fluorescence was excited with a 640 nm laser and recorded with a 670/30 nm bandpass filter and 505 nm long-pass filter. FLIPR fluorescence was excited using a 488 nm laser and recorded on a 525/50 nm bandpass filter. As before the same general sorting scheme was used to identify whole single cells based on forward and side scatter and enriched for good quality library members based on miRFP670 fluorescence. An example of the gating strategy is in **Supplemental Fig. 2** Cells were sorted based on resting membrane potential into 3 populations (miRFP^high^/FLIPR^Low^, miRFP^high^/FLIPR^medium^, miRFP^high^/FLIPR^high^. The resting membrane potential experiment was done in duplicate on separate days for two entirely independent replicates.

### Sequencing

For both biological replicates, DNA from pre-sort control and sorted cells was extracted with Microprep DNA kits (Zymo Research) and triple-eluted with water. The elute was diluted such that no more than 1.5ug of DNA was used per PCR reaction and amplified for 20 cycles of PCR using Primestar GXL (Takara Bio), run on a 1% agarose gel, and gel purified. Primers that bind outside the recombination site ensure leftover plasmid DNA from the original cell line construction step is not amplified. Purified DNA was quantified using Picogreen DNA quantification. Equal amounts (by mass) of each domain insertion sample were pooled by cell sorting category (“pre-sort control”, “surface expression”, “no surface expression”). Pooled amplicons were prepared for sequencing using the Nextera XT sample preparation workflow and sequenced using Illumina Novaseq in 2x150bp mode. Source sequencing data is available in the NCBI Sequence Raw Archive (https://www.ncbi.nlm.nih.gov/sra) under accession code PRJNA791691. Read count statistics are listed in **Supplemental Table 1A.**

### Alignment and Enrichment Calculation

Variant frequency for single mutations were determined by joining paired reads with bbmerge, aligning reads to the Kir2.1 reference gene sequence with bbmap, and identifying mutations and filtering to find the OLS programmed mutations with a custom python script. The 150 bp paired-end reads were trimmed to remove standard Illumina adapters (bbmap/resources) using BBDuk with ‘mink’ at 8 bases. Overlapping reads were corrected without merging using BBMerge with the ‘ecco’ and ‘mix’ setting. Reads were aligned to Kir2.1 sequence using BBMap using ‘maxindel’ at 500 and ‘local’ alignment. The resulting sam file was analyzed for mutations at each position and filtered to match only mutations that were programmed on the OLS chip. To calculate enrichment of mutations we used Enrich2 ^48^. Counts for each mutation were input into the Enrich2 software with weighted least squares scoring and wild type normalization. Scores were output with a standard error in a .csv file. Coverage of mapped reads is listed in **Supplemental Table 1B-C**.

### Inward Rectifier Phylogenetic Alignment and Clinvar mutation assignment

We downloaded all human inward rectifiers (KCNJ1(Kir1.1/ROMK; Uniprot P48048), KCNJ10(Kir1.2/Kir4.1; Uniprot P78508), KCNJ15(Kir1.3/Kir4.1; Uniprot Q99712), KCNJ2(Kir2.1; Uniprot P63252), KCNJ12(Kir2.2; Uniprot Q14500), KCNJ4(Kir2.3; Uniprot P48050), KCNJ14(Kir2.4; Uniprot Q9UNX9), KCNJ18(Kir2.6; Uniprot B7U540), KCNJ3(Kir3.1/GIRK1; Uniprot P48549), KCNJ6(Kir3.2/GIRK2; Uniprot P48051), KCNJ9(Kir3.3/GIRK3; Uniprot Q92806), KCNJ5(Kir3.4/GIRK4; Uniprot P48544), KCNJ16(Kir5.1; Uniprot Q9NPI9), KCNJ8(Kir6.1; Uniprot Q15842), KCNJ11(Kir6.2; Uniprot Q14654), KCNJ13(Kir7.1; Uniprot O60928) and aligned these together using MegaX ^102^. Based on this alignment, we generated a master list of Inward Rectifier numberings to translate residue numbering between Kir homologues. We assigned mutations observed Clinvar and dbSNP and their pathogenic classification (ClinVar only) to this master alignment (**Supplementary Table 2**).

## Author contributions

WCM, DN, and DS designed the work presented here. WCM and DN did the experiments with support from YH. DN processed the sequencing data. WCM and DS did the analysis and wrote the manuscript.

## Acknowledgement

We are grateful for helpful discussions with Anna Gloyn, James Fraser, Gabbriella Estevam, Eric Greene, the DMS crew, and the rest of the Fraser lab. We also thank you for taking the time to read our manuscript. This work was supported by the National Institutes of Health (1R01GM136851 to D.S.) and a University of Minnesota Genome Center Illumina S2 grant. W.C.­M. is supported by a National Science Foundation Graduate Research Fellowship and a Howard Hughes Medical Institute Gilliam Fellowship for Advanced Study.

## SUPPLEMENTARY MATERIALS FOR

**Supplementary Figure 1:**
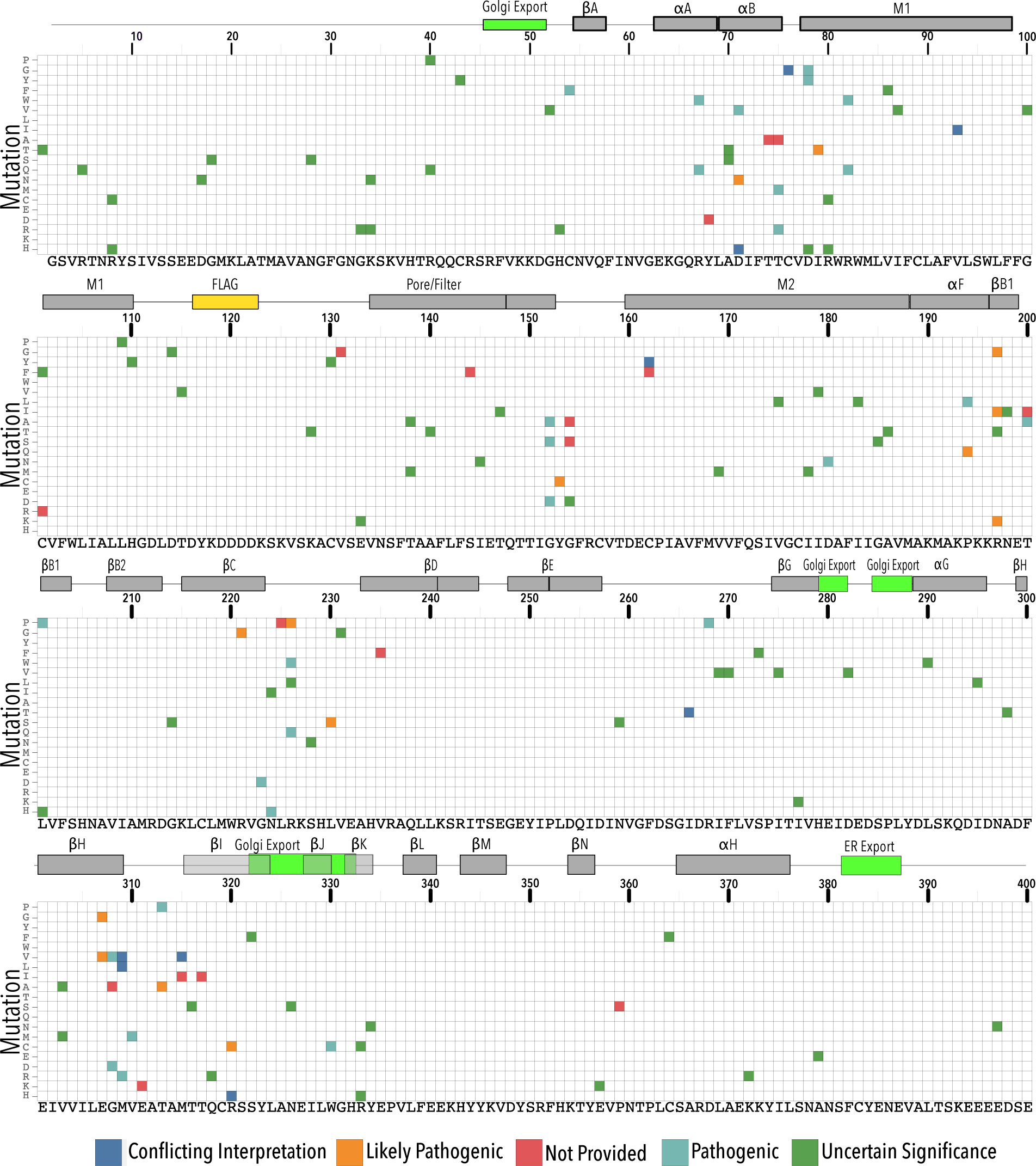
Kir2.1 variant effect reported in ClinVar. Heatmap showing clinically observed Kir2.1 variants mapped by position, mutation, and pathogenicity interpretation. Kir2.1 secondary structural element are shown as gray boxes above each panel. Trafficking signals are colored green.

**Supplementary Figure 2:**
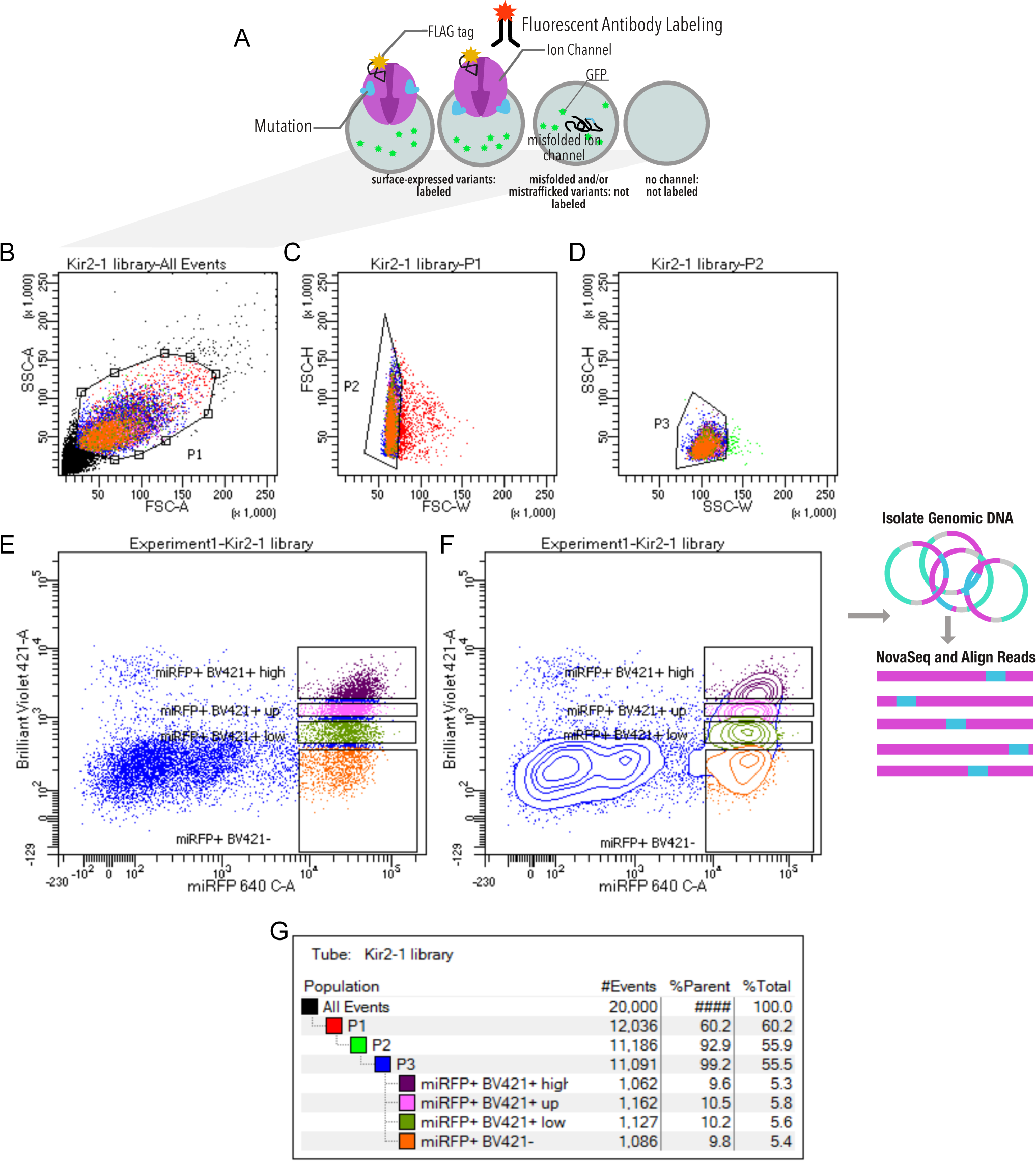
Kir2.1 surface expression assay gating scheme. **(A)** HEK293 cells expressing a singie variant are incubated with a fluorescent (Brilliant Violet) antibody that recognizes a FLAG tag inserted into an extracellular loop (Kir2.1 position T115). Cells expressing channel variants that can fold, assemble and traffic to the cell surface are labelled, while those that express variants with impaired folding, assembly, and/or trafficking are not labelled. **(B)** Using FACS, whole HEK293 cells are gated on side (SSC-A) and forward scattering (FSC-A). **(C-D)** Forward scattering height (FSC-H), forward scattering width (FSC-W), and Side scattering width (SSC-W) are used to gate single cells. Non-surface expressed variants (miRFPhigh/Labeliow) and surface expressed variants (miRFPhigh/Labelhigh) populations are gated into ’negative^1^, ’low^1^, ’up’, and ’high’ populations based an expression marker (miRFP670) of Anti-Flag Brilliant Violet-421 fluorescence. Scatter plot **(E)** and contour plot **(F)** shown. Contour plots represent 95% confidence intervals with outliers shown as dots. **(G)** Sort statistic for each gated cell population.

**Supplementary Figure 3:**
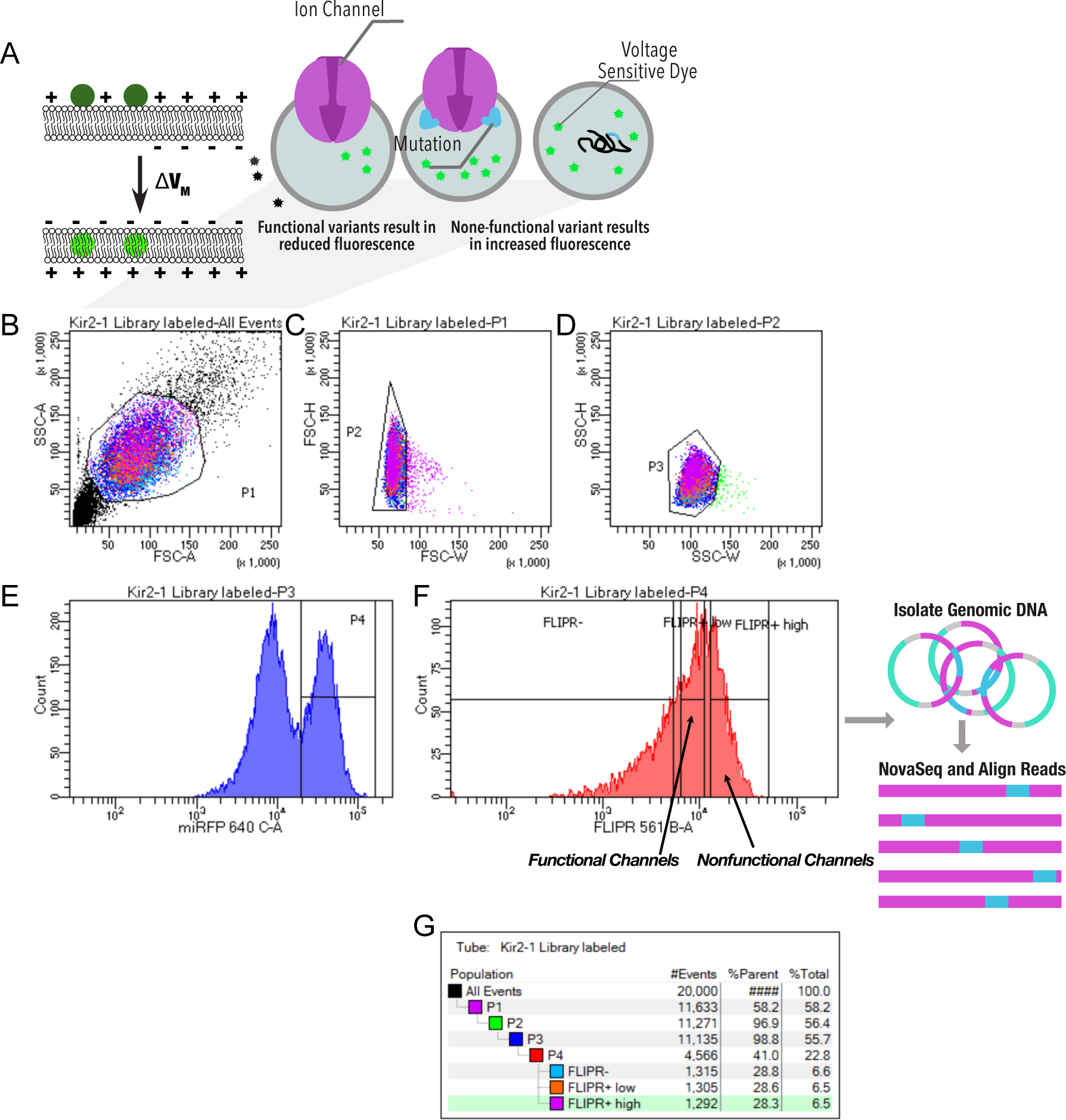
Kir2.1 functional fitness assay gating scheme. **(A)** HEK293 cells expressing a single variant are incubated with a voltage-sensitive FLIPR dye. In strongly hyperpolarized cells, this dye partitions out the cell membrane into an aqueous environment, which decreases its extinction coefficient and thus decreases green fluorescence upon excitation with blue light. In depolarized cells, the FLIPR dye fluorescence is brighter because the dye partitions into the membrane, which increases its extinction coefficient. **(B)** Using FACS, whole HEK293 cells are gated on side (SSC-A) and forward scattering (FSC-A). **(C-D)** Forward scattering height (FSC-H), forward scattering width (FSC-W), and Side scattering width (SSC-W) are used to gate single cells. **(E)** Cells are further gated using miRFP670 as an expression marker. **(F)** Non-functional and functional channel population are separated by sorting based on FLIPR fluorescences. **(G)** Sort statistic for each gated cell population.

**Supplementary Figure 4:**
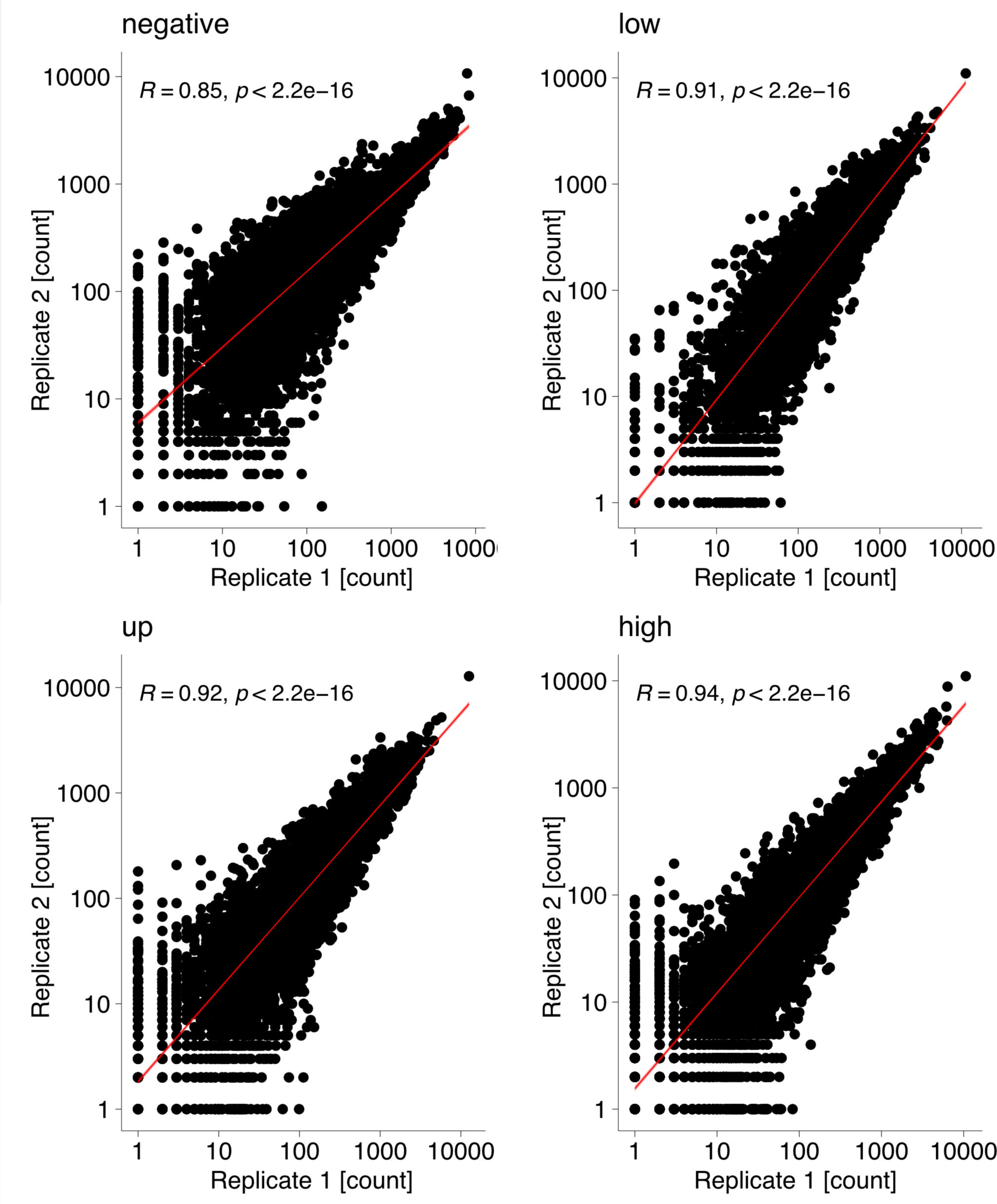
Surface fitness assay biological replicates. Scatterplots show mapped read counts for each Kir2.1 variant in each replicate for different assay subpools. Spearman correlation coefficients are inset for each replicate pair. A linear model regression fit is shown as a red line.

**Supplementary Figure 5:**
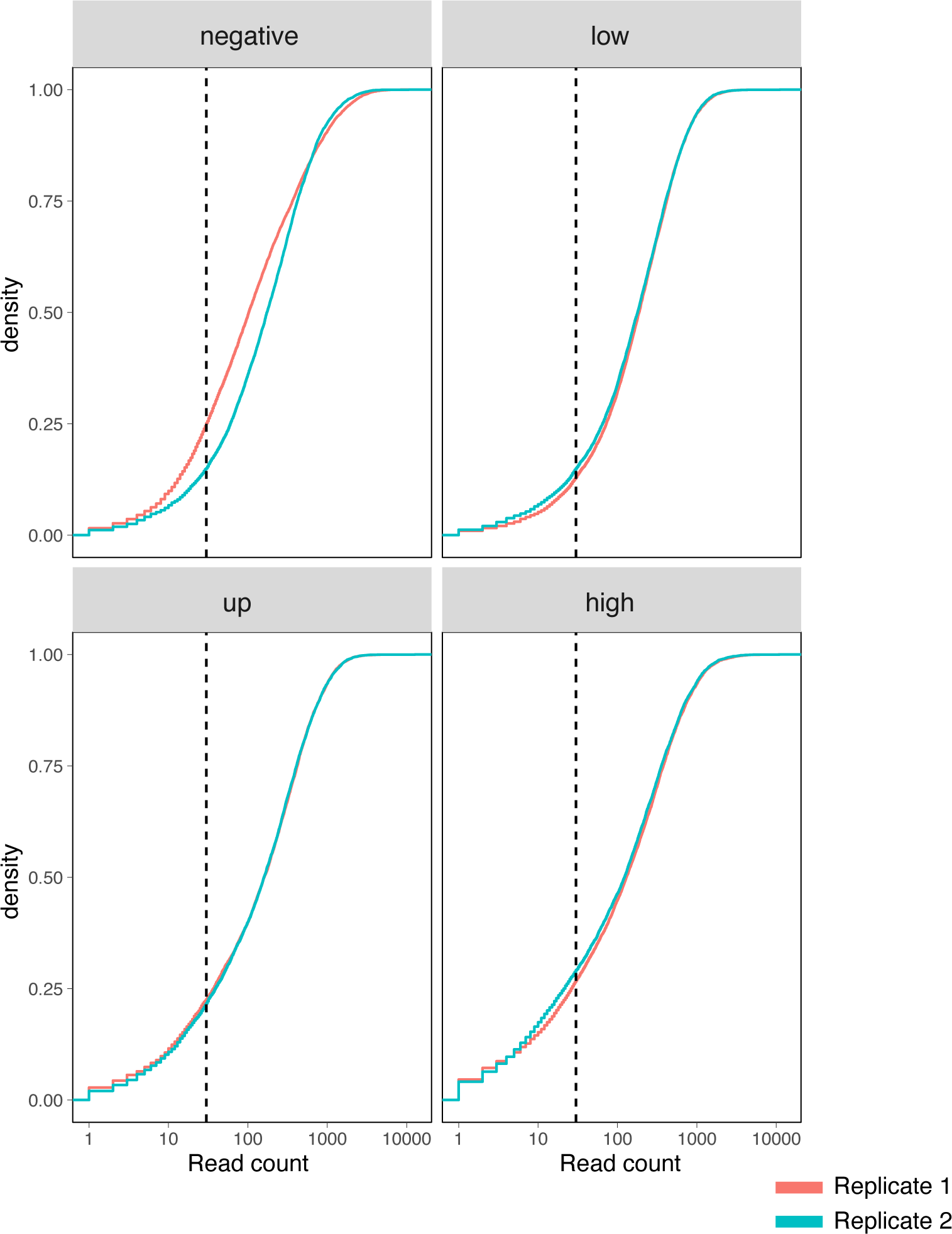
Surface fitness assay read count depth. Cumulative density distribution of read counts for each replicate .and assay condition subpool. Dashed vertical lines represent 30-fold coverage.

**Supplementary Figure 6:**
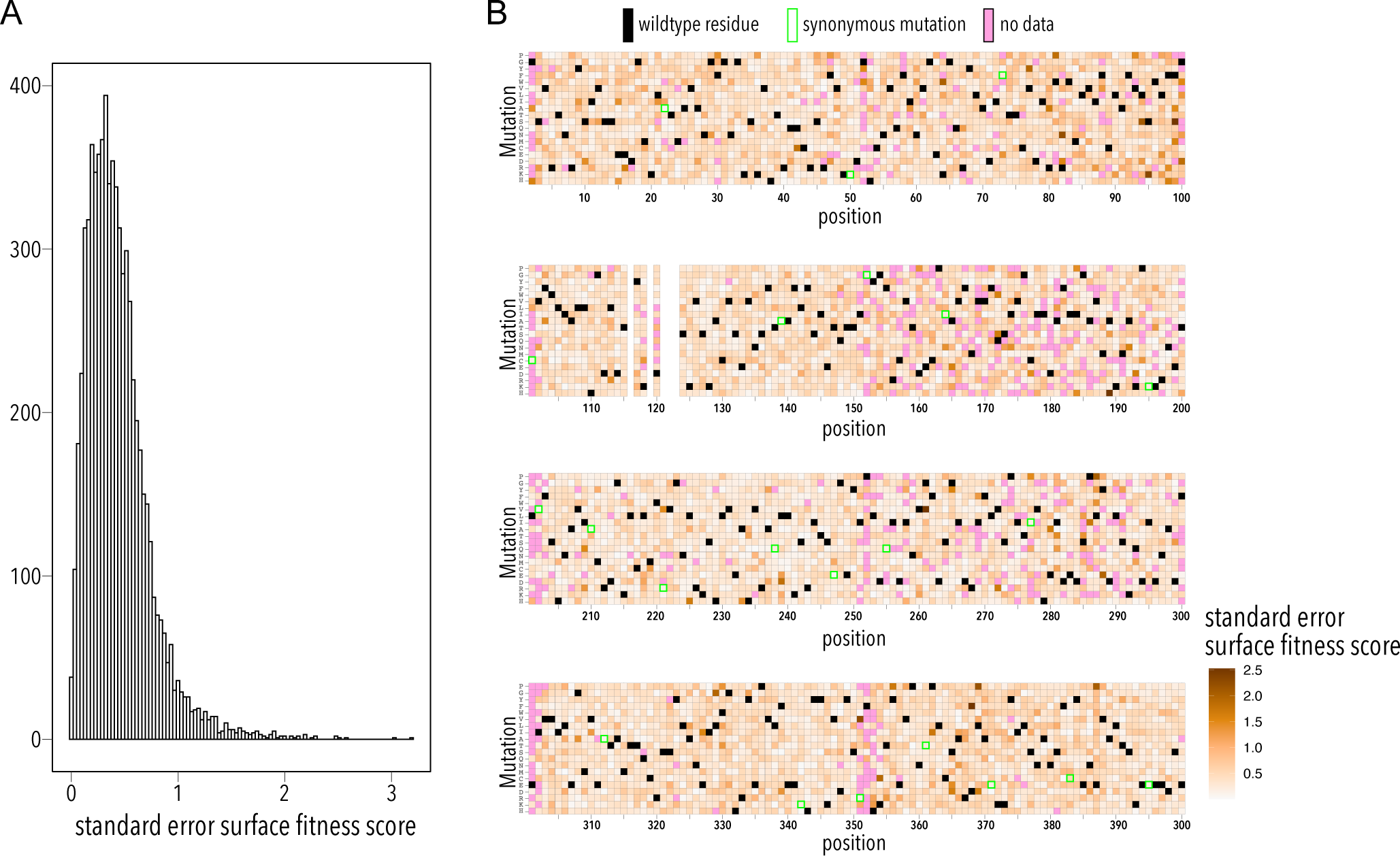
Surface fitness assay standard error distribution. **(A)** Distribution of surface fitness score standard errors. **(B)** Heatmap showing surface fitness standard error (gradient white to brown) mapped to each position and mutation. Wildtype residues are shown as black boxes. Missing data is shown as magenta boxes. Synonymous mutations are indicated by a green outline.

**Supplementary Figure 7:**
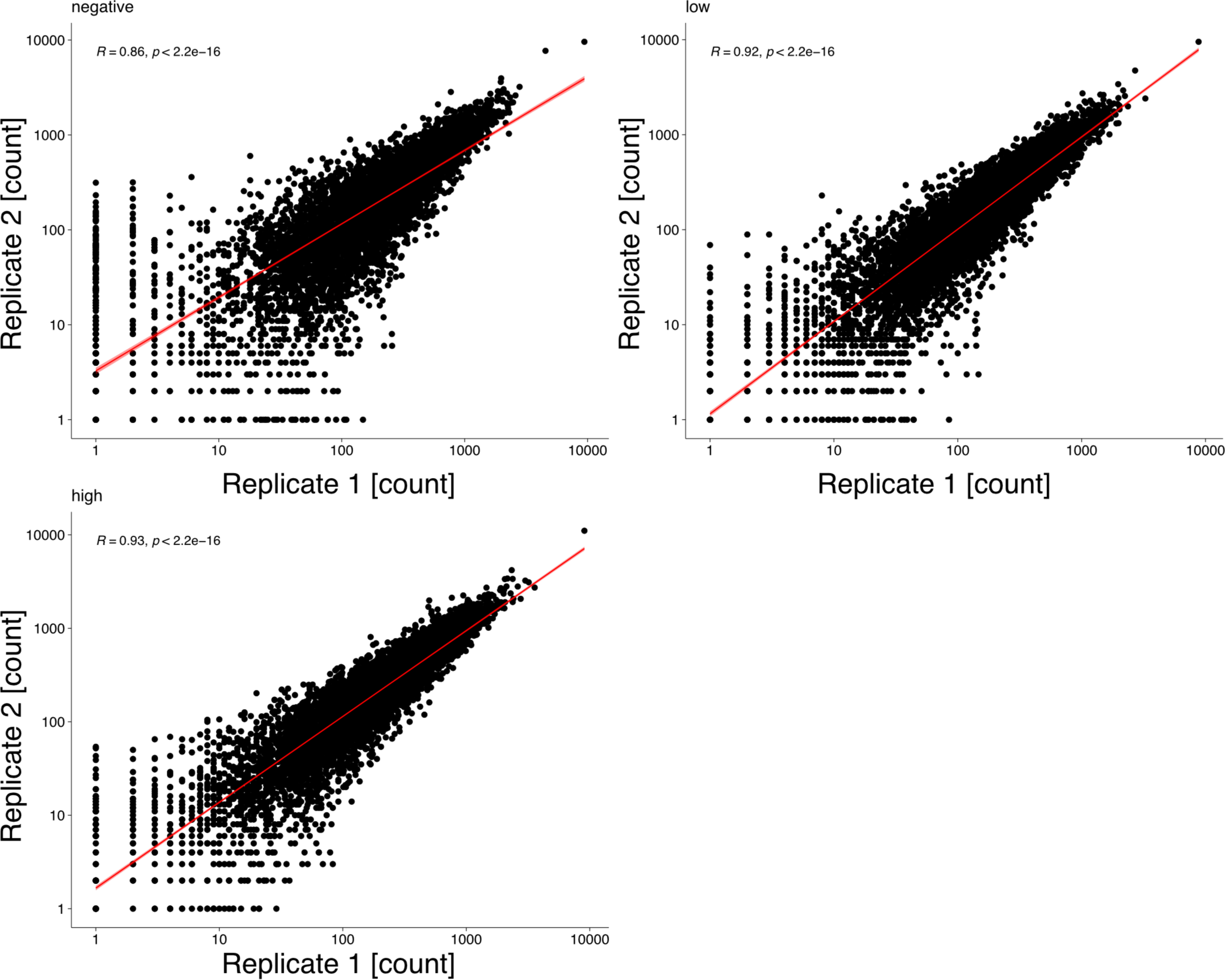
Surface fitness assay biological replicates. Scatterplots show mapped read counts for each Kir2.1 variant in each replicate for different assay subpools. Spearman correlation coefficients are inset for each replicate pair. A linear model regression fit is shown as a red line.

**Supplementary Figure 8:**
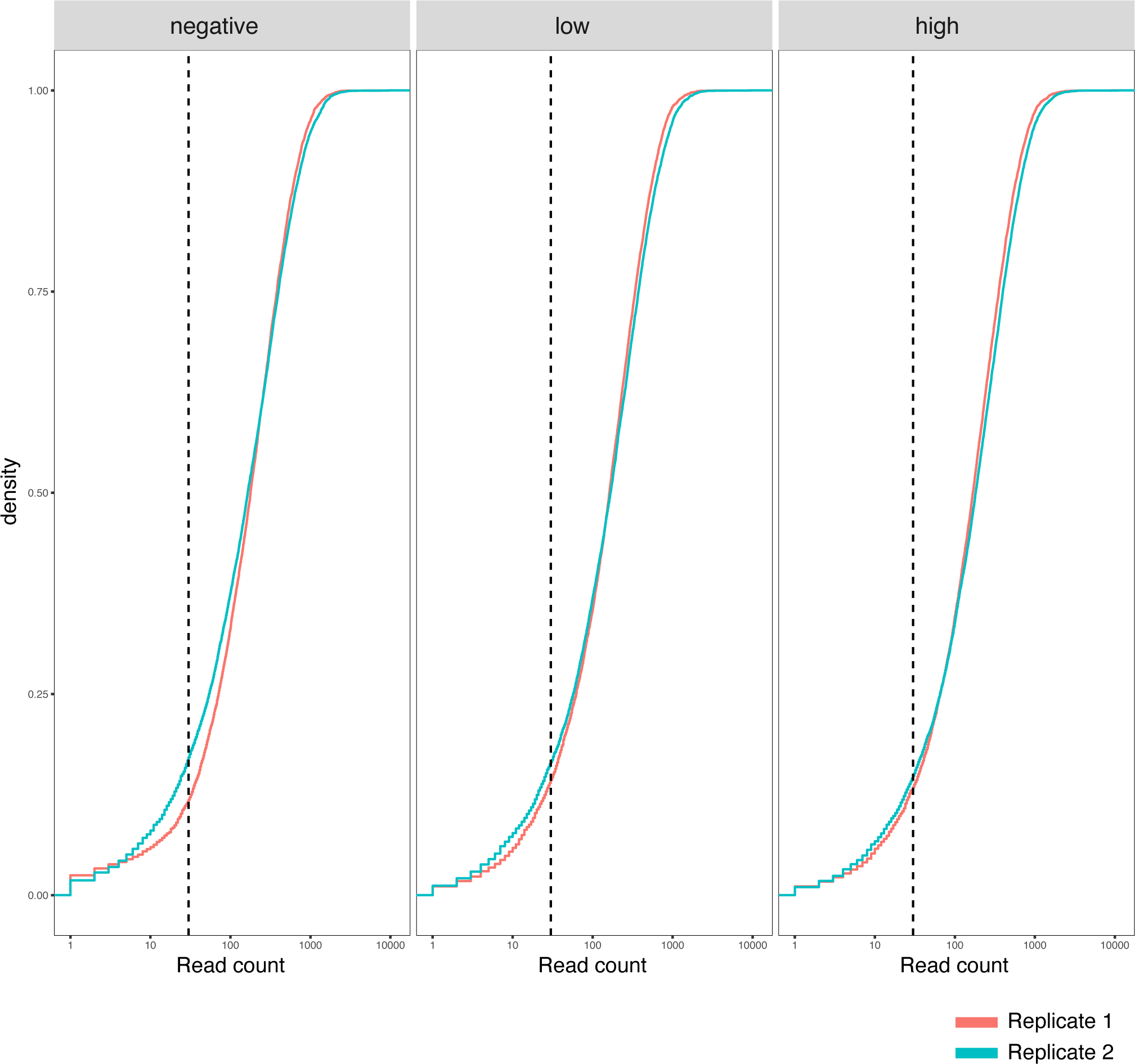
Surface fitness assay biological replicates. Cumulative density distribution of read counts for each replicate and assay condition subpool. Dashed vertical lines represent 30-fold coverage.

**Supplementary Figure 9:**
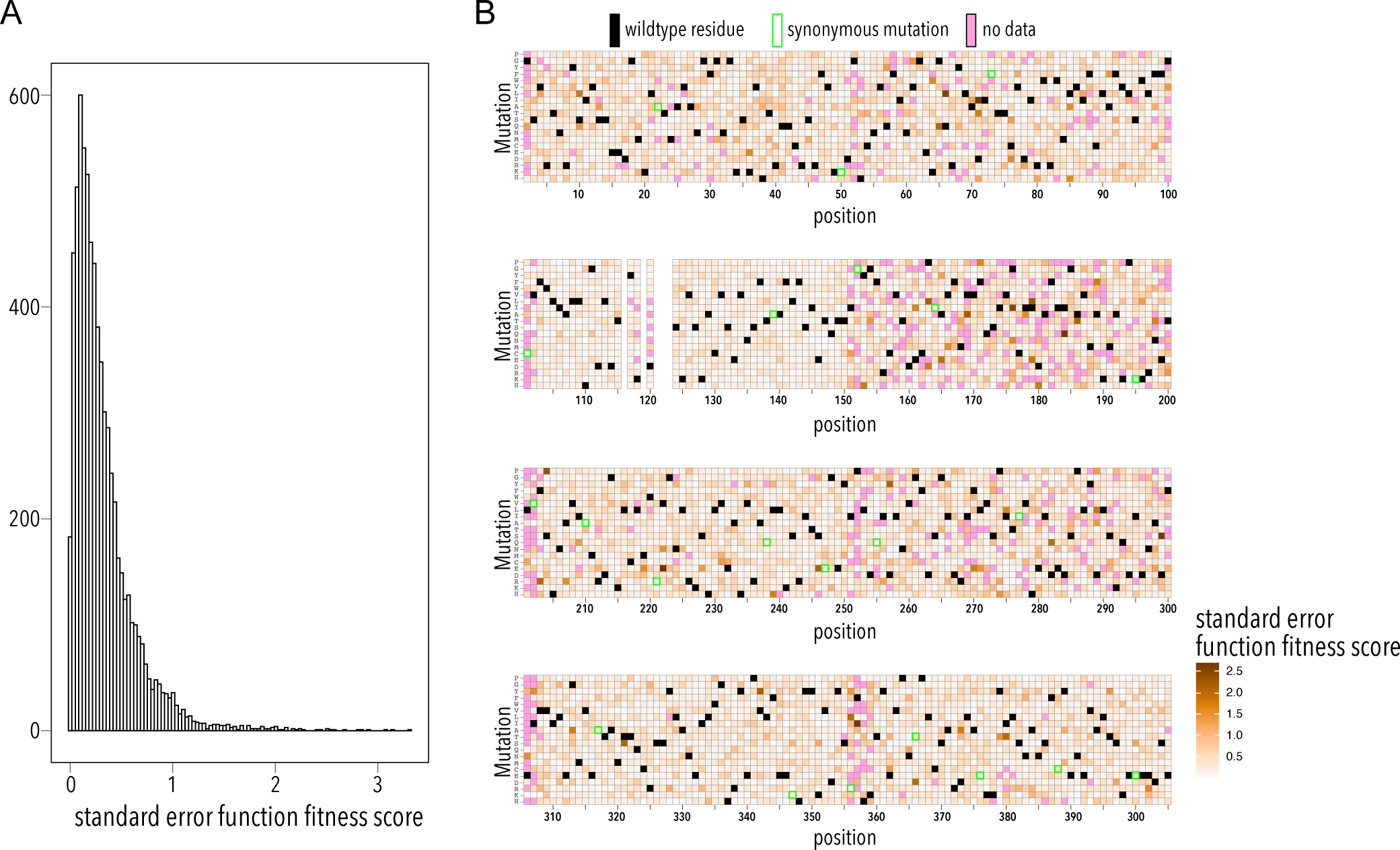
Function fitness assay standard error distribution. **(A)** Distribution of surface fitness score standard errors. **(B)** Heatmap showing surface fitness standard error (gradient white to brown) mapped to each position and mutation. Wildtype residues are shown as black boxes. Missing data is shown as magenta boxes. Synonymous mutations are indicated by a green outline.

**Supplementary Table 1:** NGS read statistics and assay repeatability. **(A)** Read statistics tor each sequenced variant pool. Mapped reads for each replicate and surface fitness assay sub-pool **(B)** or function fitness assay **(C). Nvariants** represented in each replicate (out of 7,429 total), median read count per variant, mean read count per variant, and percentage of variants that have greater than 30-fold reach coverage.

| Sample | Barcode sequence | PF Clusters | % of the lane | % Perfect barcode | % One mismatch barcode | Yield (Mbases) | % PF Clusters | % ≥Q30 bases | Mean Quality Score |
| --- | --- | --- | --- | --- | --- | --- | --- | --- | --- |
| dms_baseline | CGTACTAG+AGAGGATA | 17,942,041 | 1.78 | 95.35 | 4.65 | 5,418 | 100 | 90.5 | 35.34 |
| dms_function_high1 | GTAGAGGA+ATAGAGAG | 14,044,823 | 1.39 | 95.15 | 4.85 | 4,242 | 100 | 87.22 | 34.64 |
| dms_function_high2 | TAAGGCGA+AGAGGATA | 18,810,378 | 1.86 | 96.5 | 3.5 | 5,681 | 100 | 89.08 | 35.01 |
| dms_function_low1 | CGAGGCTG+ATAGAGAG | 14,199,546 | 1.41 | 97.46 | 2.54 | 4,288 | 100 | 88.63 | 34.99 |
| dms_function_low2 | GGAGCTAC+ATAGAGAG | 14,903,201 | 1.48 | 95.74 | 4.26 | 4,501 | 100 | 89.28 | 35.1 |
| dms_function_neg1 | TAGGCATG+ATAGAGAG | 15,073,953 | 1.49 | 97.14 | 2.86 | 4,552 | 100 | 91.1 | 35.45 |
| dms_function_neg2 | ACTCGCTA+ATAGAGAG | 16,560,022 | 1.64 | 95.44 | 4.56 | 5,001 | 100 | 89.56 | 35.14 |
| dms_surface_high1 | TCCTGAGC+ATAGAGAG | 15,200,690 | 1.51 | 96.22 | 3.78 | 4,591 | 100 | 90.8 | 35.37 |
| dms_surface_high2 | TAGGCATG+AGAGGATA | 14,882,582 | 1.47 | 97.33 | 2.67 | 4,495 | 100 | 90.87 | 35.4 |
| dms_surface_low1 | CGTACTAG+ATAGAGAG | 16,320,952 | 1.62 | 93.92 | 6.08 | 4,929 | 100 | 88.75 | 34.98 |
| dms_surface_low2 | TCCTGAGC+AGAGGATA | 17,274,356 | 1.71 | 97.2 | 2.8 | 5,217 | 100 | 90.98 | 35.42 |
| dms_surface_neg1 | TAAGGCGA+ATAGAGAG | 18,343,213 | 1.82 | 94.16 | 5.84 | 5,540 | 100 | 87.59 | 34.73 |
| dms_surface_neg2 | AGGCAGAA+AGAGGATA | 20,844,383 | 2.07 | 96.24 | 3.76 | 6,295 | 100 | 90.49 | 35.31 |
| dms_surface_up1 | AGGCAGAA+ATAGAGAG | 16,105,434 | 1.6 | 94.92 | 5.08 | 4,864 | 100 | 89.01 | 35.01 |
| dms_surface_up2 | GGACTCCT+AGAGGATA | 16,752,187 | 1.66 | 97.54 | 2.46 | 5,059 | 100 | 90.65 | 35.36 |

Surface Fitness Assay – Mapped Reads
|  | Negative |  | Low |  | Up |  | High |  |
| --- | --- | --- | --- | --- | --- | --- | --- | --- |
|  | Rep. 1 | Rep. 2 | Rep. 1 | Rep. 2 | Rep. 1 | Rep. 2 | Rep. 1 | Rep. 2 |
| $N_{\text{variants}}$ | 7,282 | 7,282 | 7,305 | 7,305 | 7,175 | 7,175 | 6,975 | 6,975 |
| median | 103 | 174 | 187 | 179 | 160.5 | 160 | 130 | 123 |
| mean | 342.3 | 344.6 | 315.5 | 307.4 | 310.4 | 309.5 | 301.4 | 285.5 |
| >30-fold [%] | 71.7 | 82.4 | 85.4 | 82.2 | 73.4 | 73.4 | 65.7 | 64.4 |

|  | Negative |  | Low |  | Up |  |
| --- | --- | --- | --- | --- | --- | --- |
|  | Rep. 1 | Rep. 2 | Rep. 1 | Rep. 2 | Rep. 1 | Rep. 2 |
| N | 7,168 | 7,168 | 7,284 | 7,284 | 7,320 | 7,320 |
| median | 178 | 165 | 162 | 167 | 168 | 182 |
| mean | 281.6 | 297.1 | 248.5 | 276.5 | 261.5 | 295.7 |
| >30-fold [%] | 79.4 | 79.5 | 82.7 | 80.9 | 83.3 | 83.5 |

**Supplemental Table 2:**
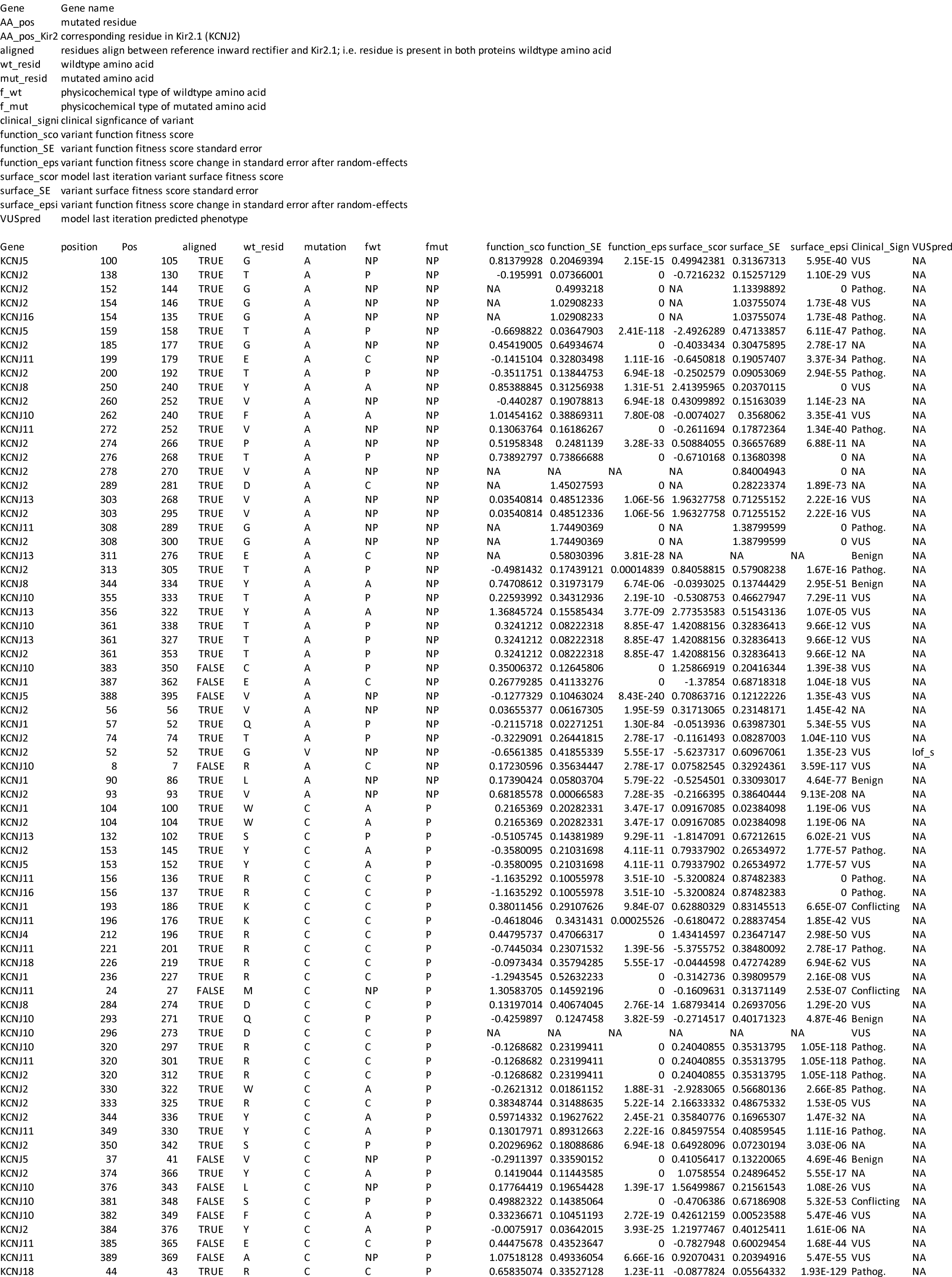
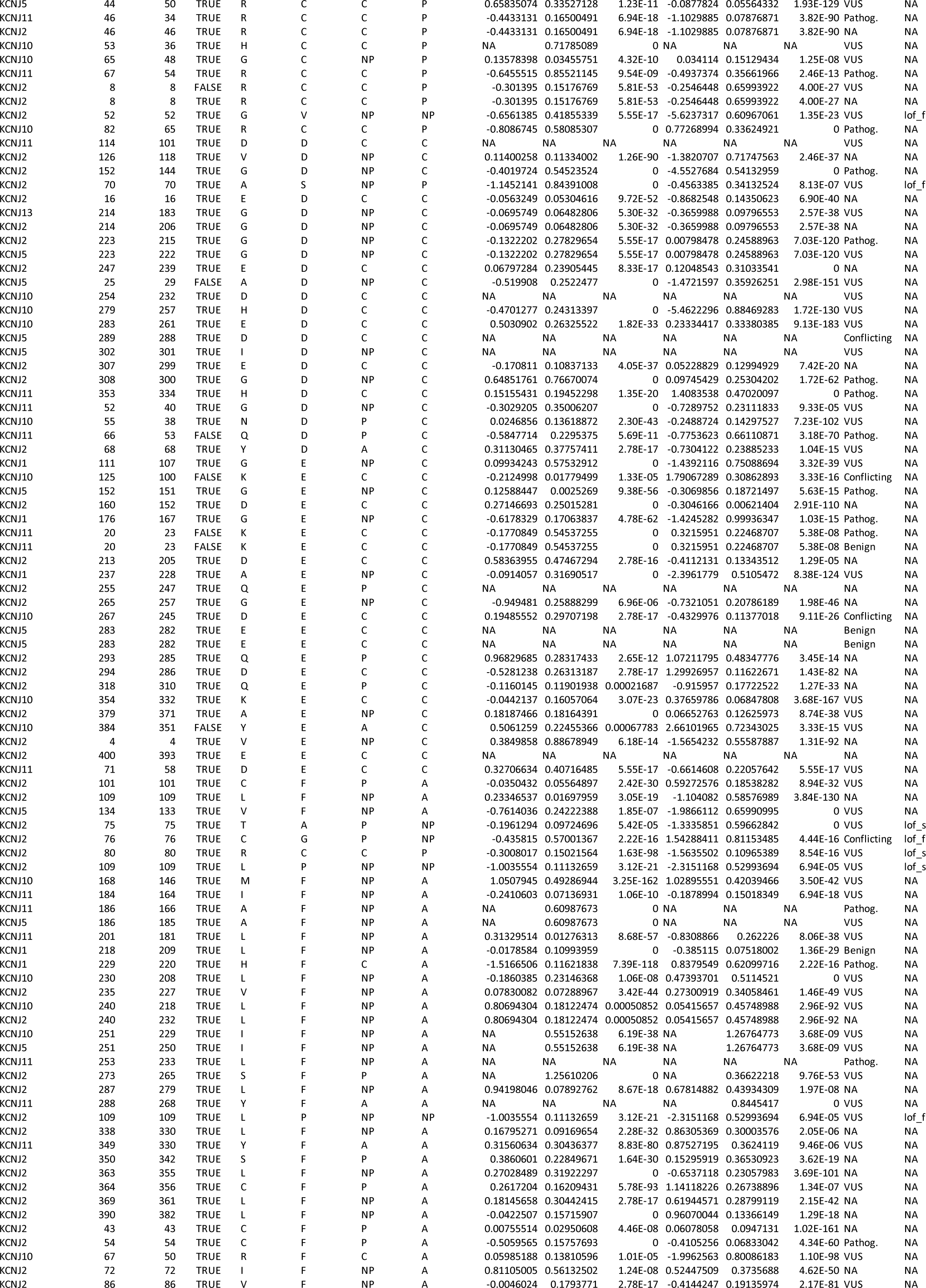
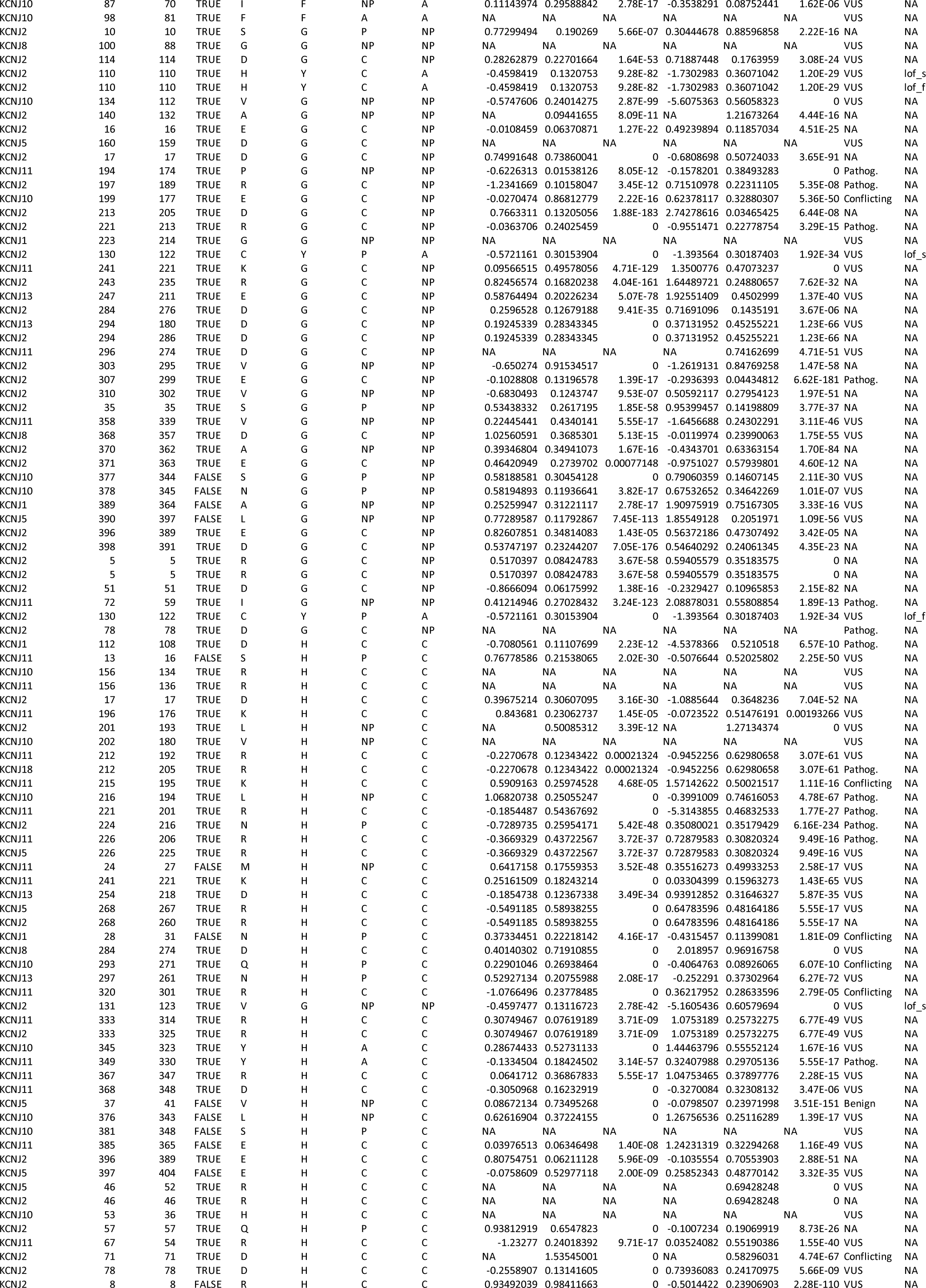
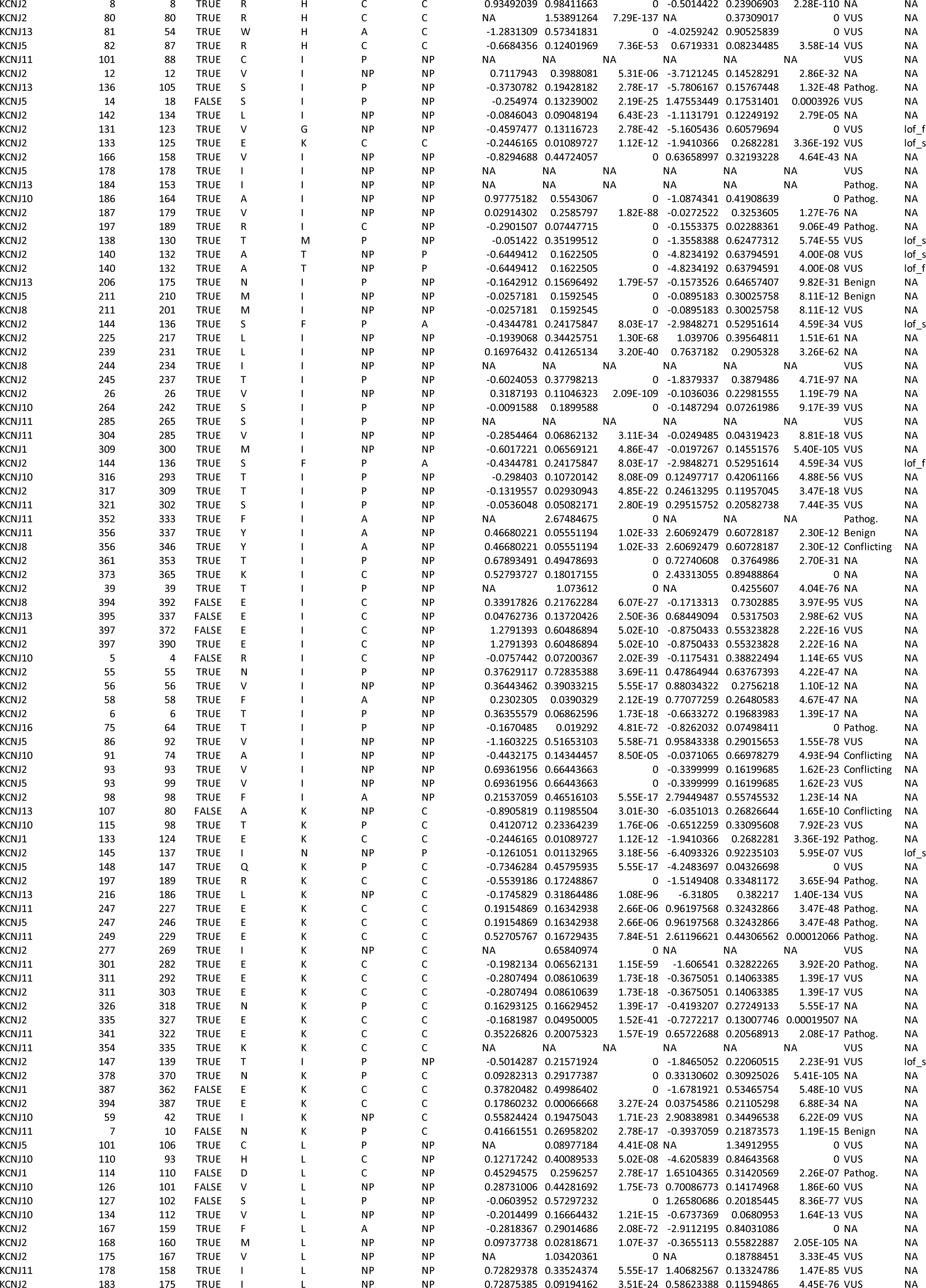
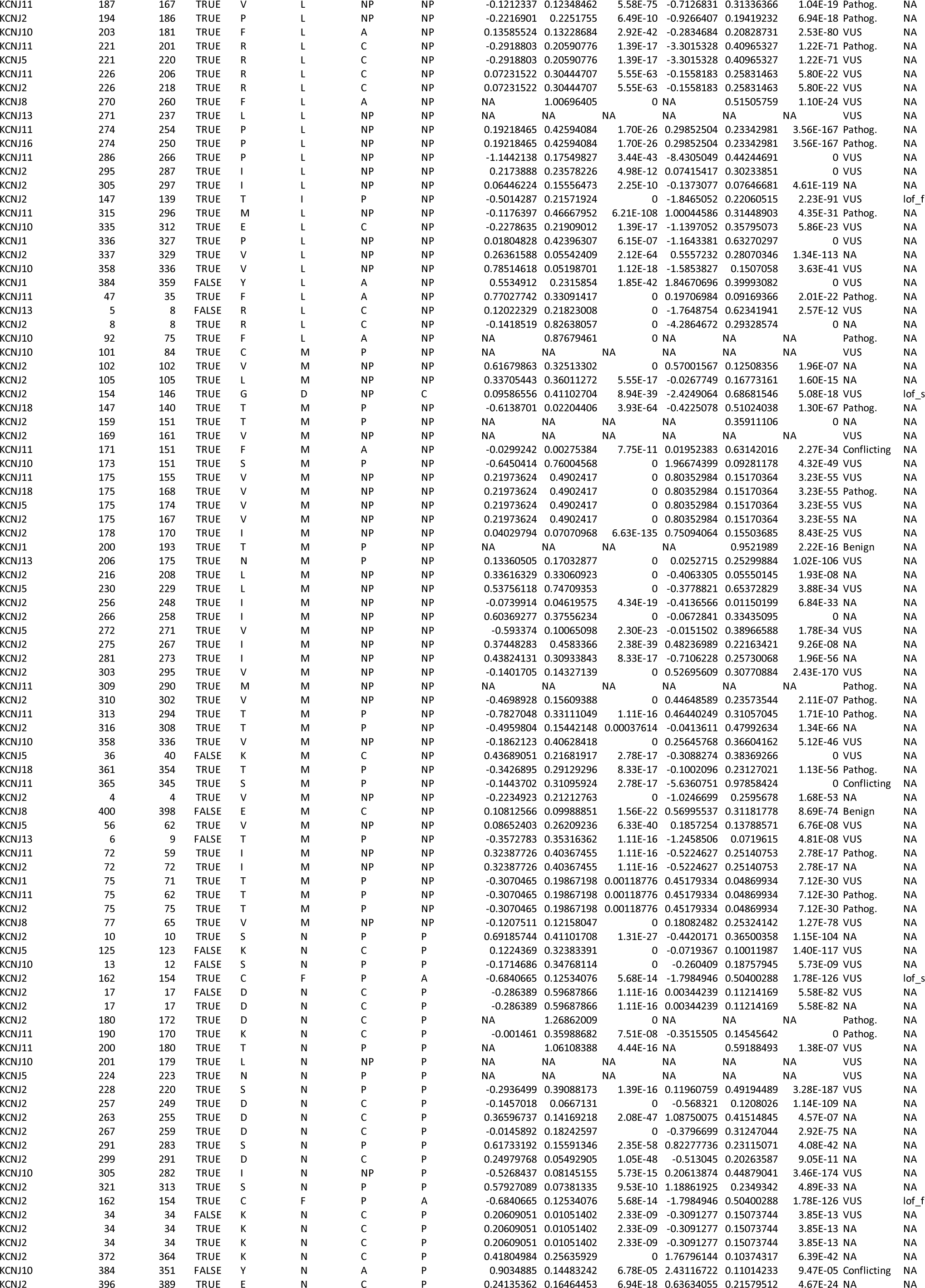
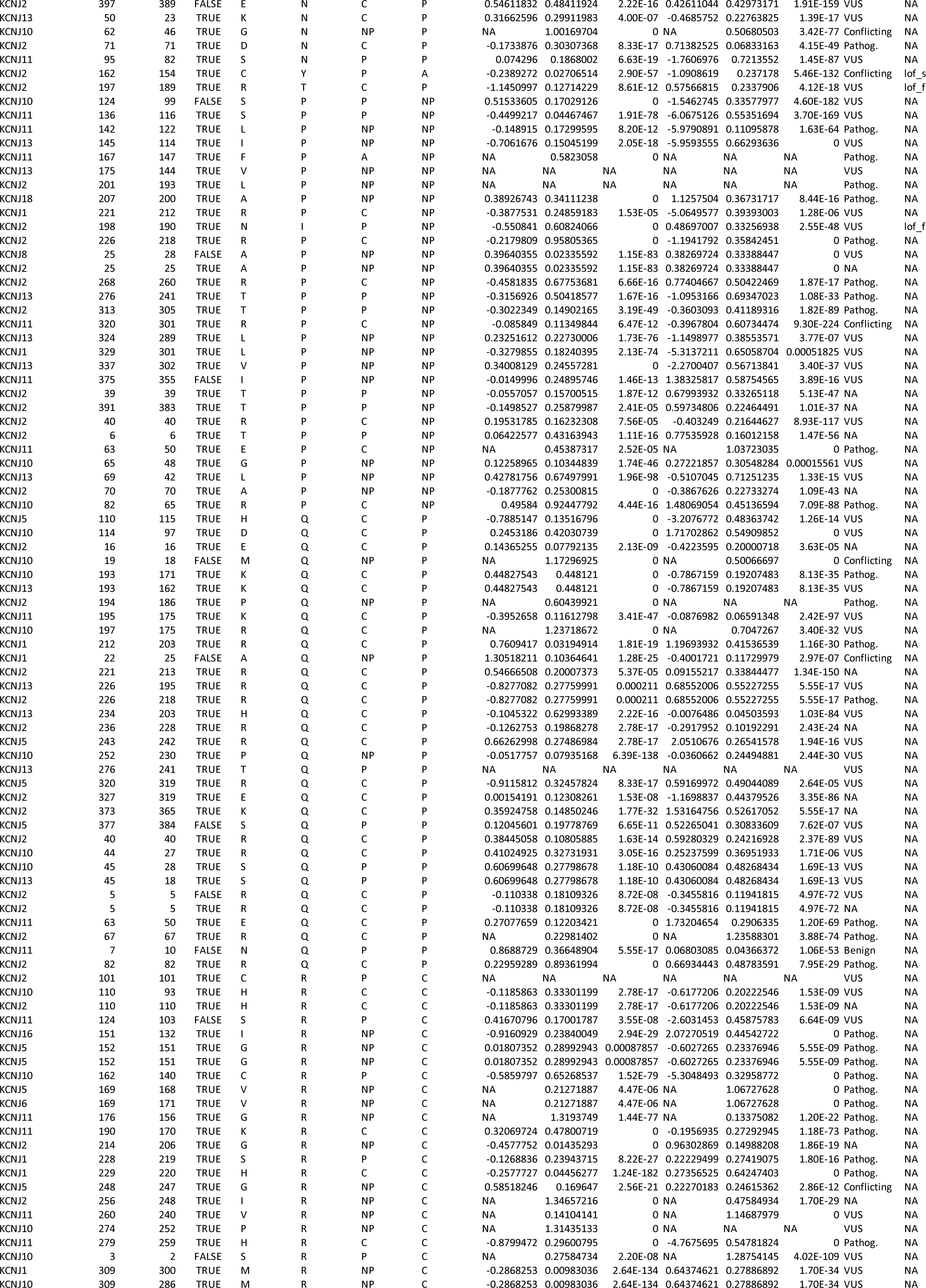
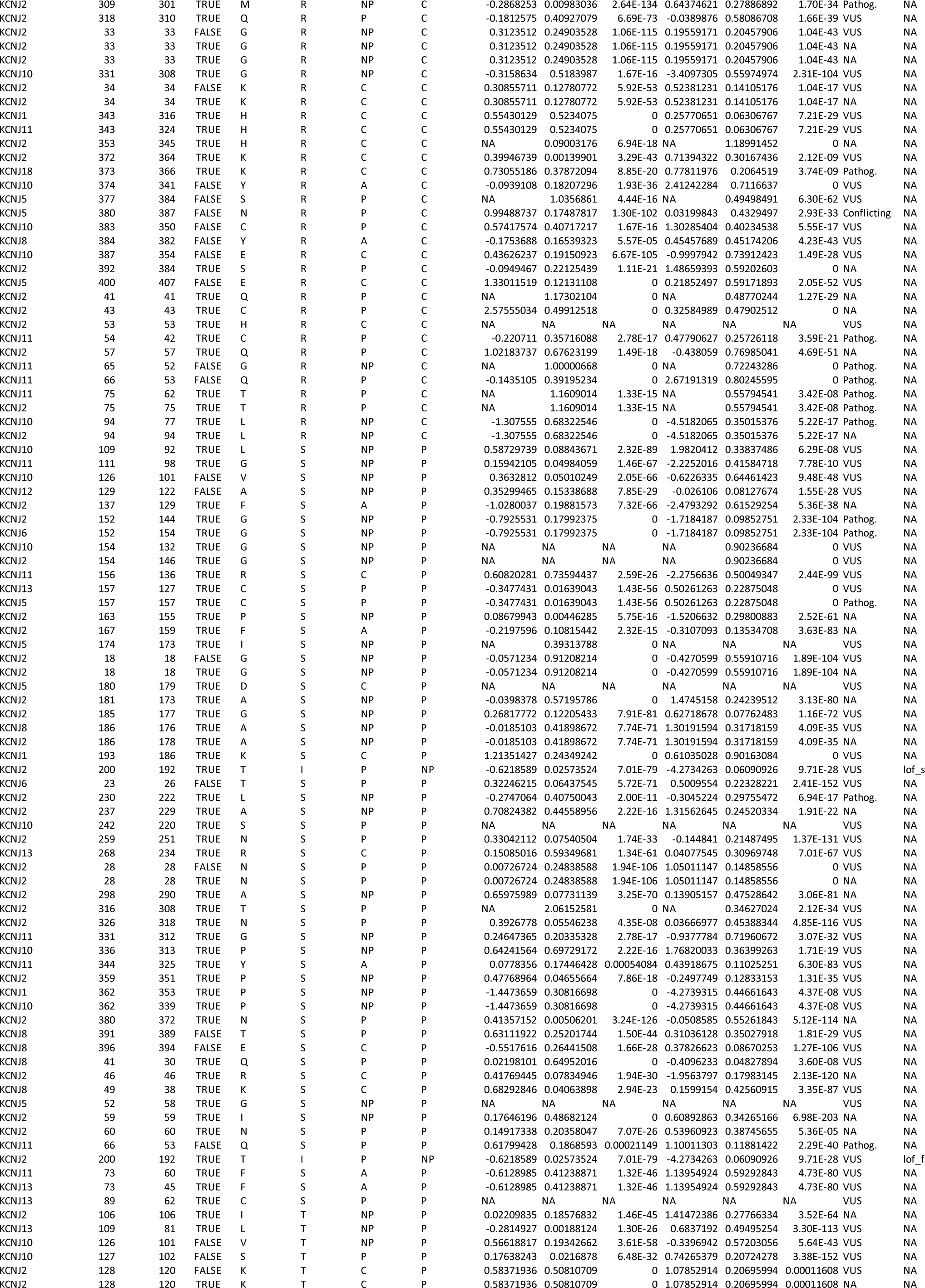
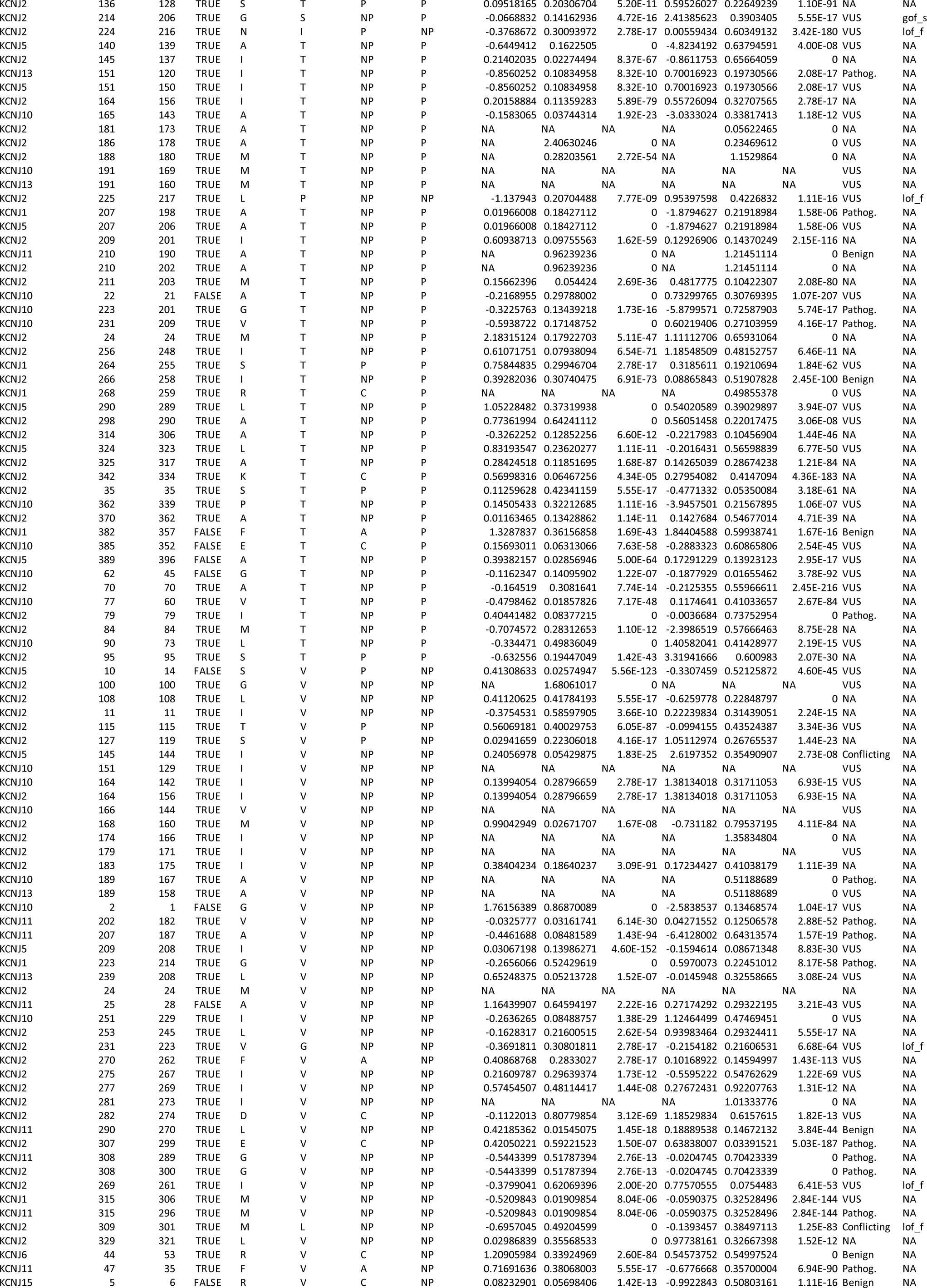
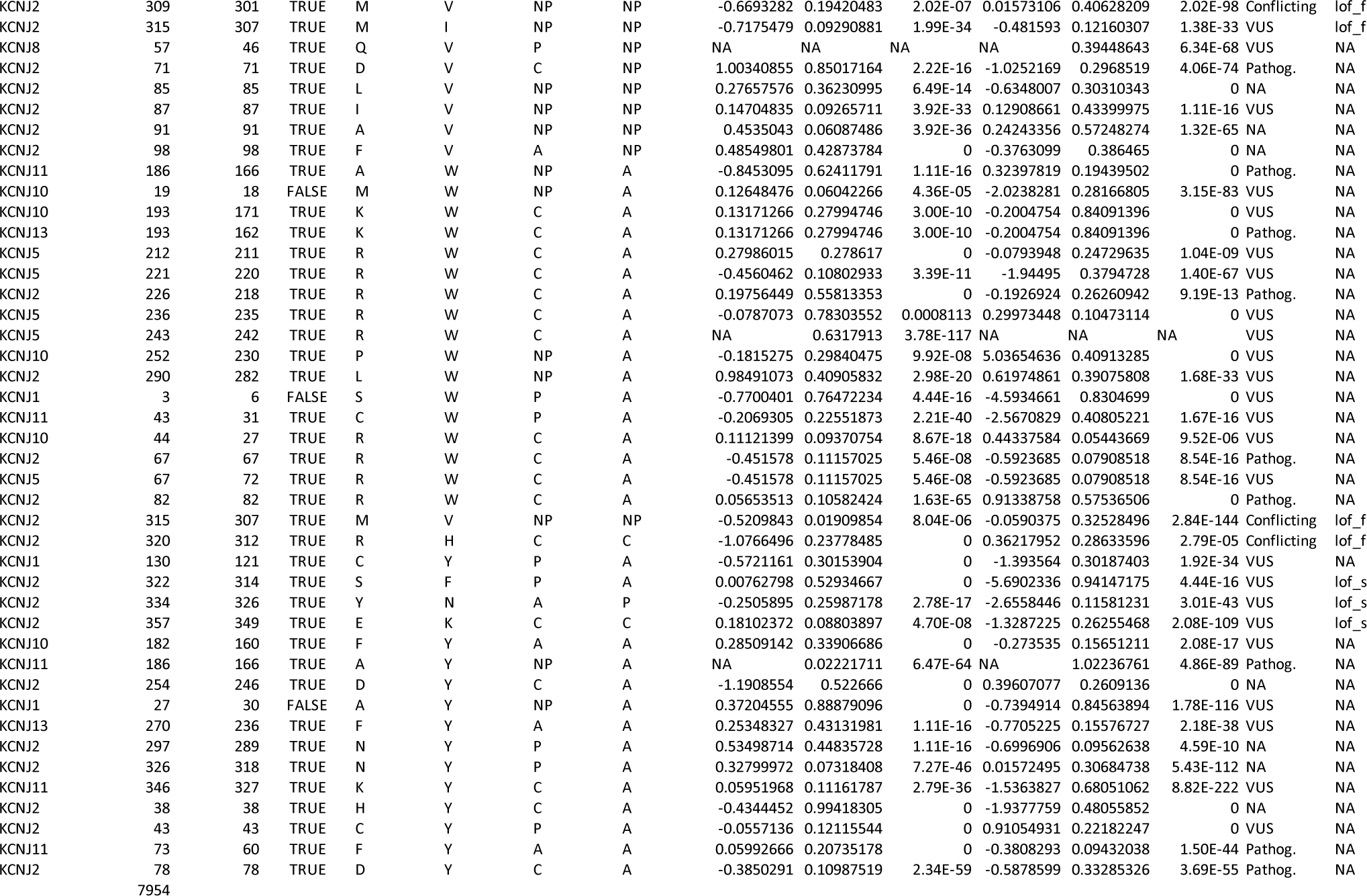
Inward Rectifier Missense Variants in ClinVar and dbSNP

**Supplemental Table 3:**
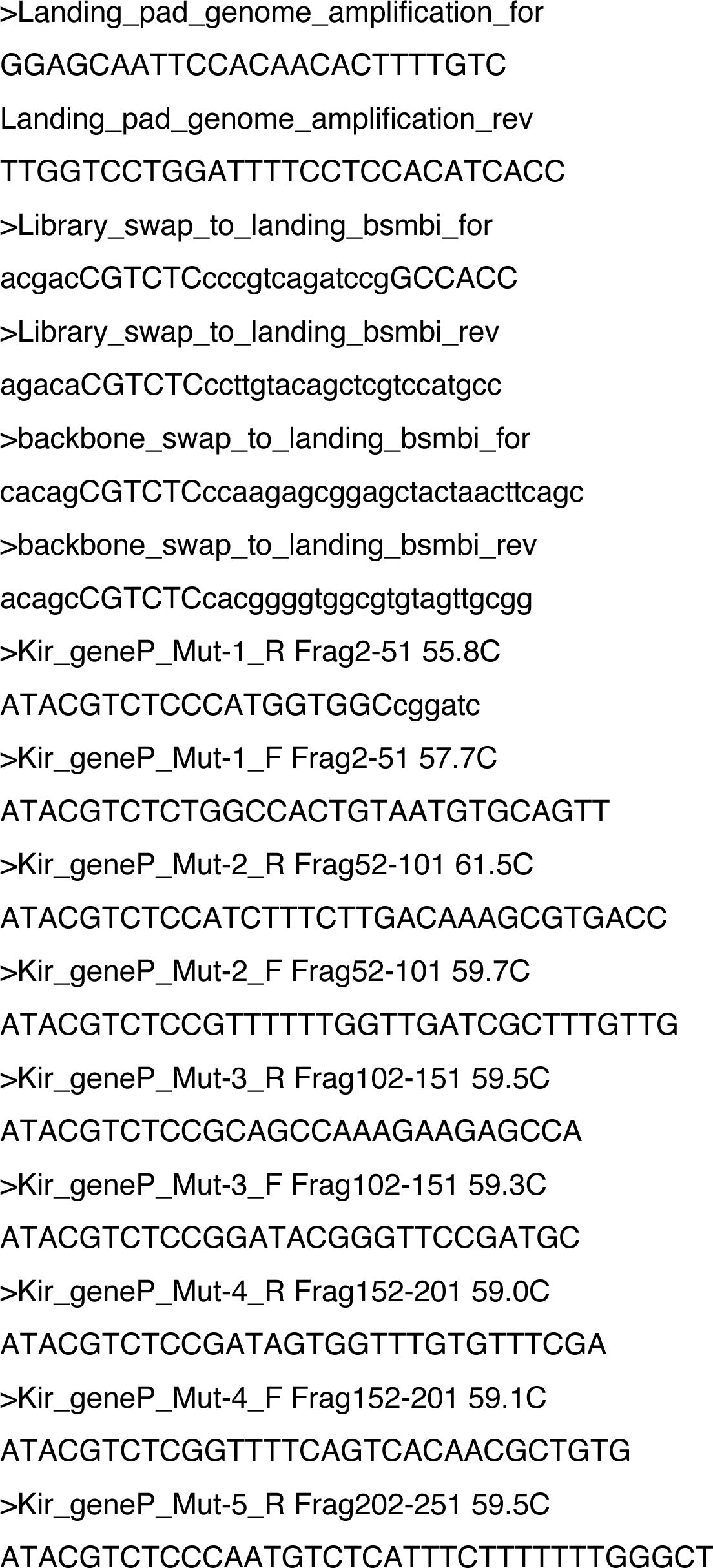
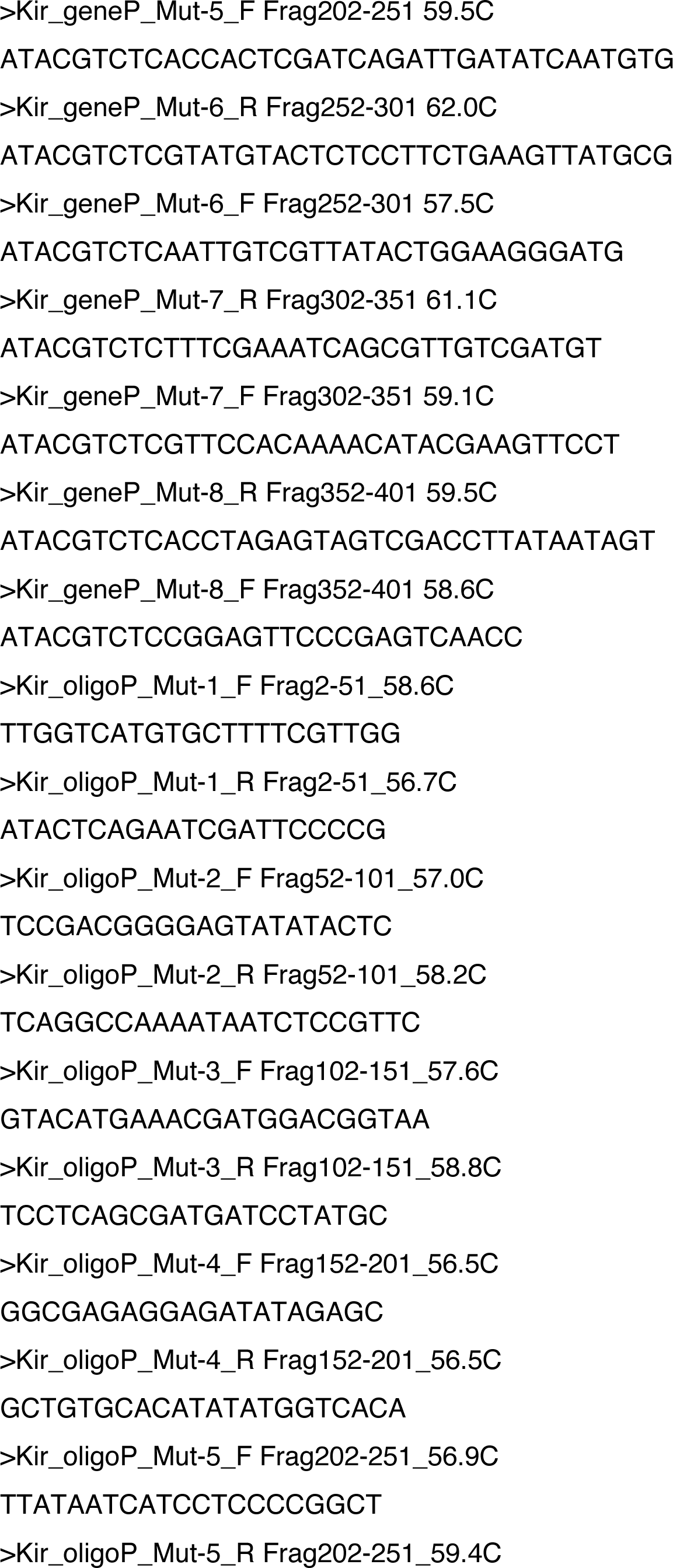
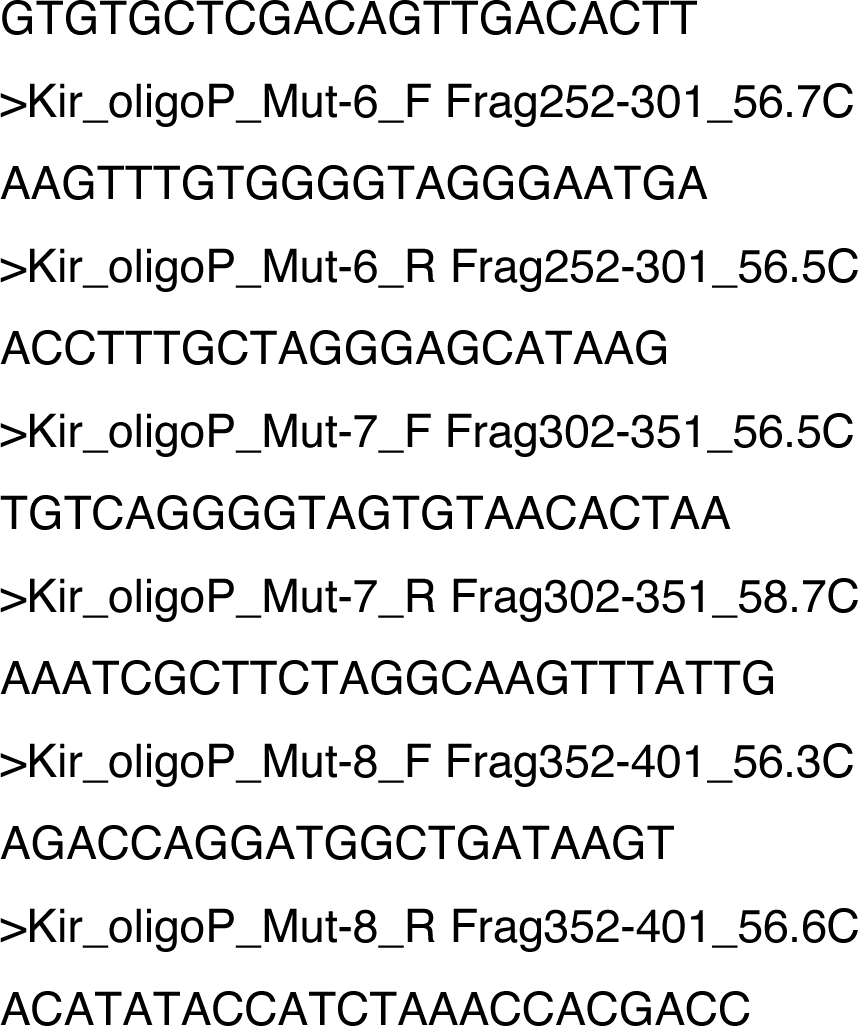
Primers used in study.

